# Decoding human B cell ontogeny in prenatal and adult bone marrow and in vitro models via single-cell multiomics

**DOI:** 10.1101/2025.08.31.672613

**Authors:** Roser Vilarrasa-Blasi, Seungjoon Kim, J. Patrick Pett, Jakov Korzhenevich, Michael Svaton, Chenqu Suo, Frederick C.K. Wong, Oxana Nashchekina, Di Zhou, Ryan Colligan, Yale S. Michaels, Carla Zimmerman, Kevin Troule, Hananeh Aliee, Valentina Lorenzi, Marie Moullet, Carmen Sancho-Serra, Davide Carra, Leonardo Morsut, Elena Prigmore, Martin Prete, Iva Kelava, Sarah A. Teichmann, Klaus Warnatz, Biola M. Javierre, Luz Garcia-Alonso, Peter W. Zandstra, Marta Rizzi, Roser Vento-Tormo

## Abstract

In humans, the bone marrow becomes the primary site for B lymphopoiesis during the second trimester of pregnancy and continues throughout life. Prenatal and adult B cell progenitors play distinct roles in the aetiology and pathology of paediatric and adult hematopoietic malignancies, though the molecular drivers of these differences remain unclear. Here, we created a comprehensive multiomics atlas of over 500k cells covering immune and stromal compartment from prenatal and adult bone marrow, enabling high-resolution analysis of the cell-intrinsic and cell-extrinsic processes that modulate prenatal and adult B lymphopoiesis. Even though B cells follow broadly similar developmental trajectories, we identify a novel postnatal ‘lateCLP’ subset equivalent to prenatal ‘preProB’ cells, and uncover prenatal cells carry several signatures characteristic of leukemias, including enhanced proliferation, higher *RAG1/RAG2* activity, extrinsic B cell signals such as *IL7*, and lower retention signals in the bone marrow. We also developed and characterised, at both cellular and molecular levels, a new experimental framework for generating B cell precursors from human induced pluripotent stem cells (hiPSCs), and show that it faithfully recapitulates key stages of B cell differentiation. Together, our single-cell multiomics atlas of B lymphopoiesis *in vivo* and *in vitro* offers detailed insights into the unique molecular features of prenatal and adult B cell lymphopoiesis, and serves as a powerful resource for investigating the early events that contribute to haematological disorders.

## Introduction

B lymphopoiesis involves multiple coordinated steps that ensure the continuous generation of immature B cells, which populate the periphery. This process is tightly regulated through key mechanisms including coordinated expression of fate determining transcription factors (TFs)^1^, stepwise rearrangement of the immunoglobulin receptor^2^, stage specific proliferation^3^, signals from the microenvironment^4,5^ and negative selection against self-reactive specificities^6^. In humans, fetal hematopoietic stem cells (HSCs) initiate commitment to the B cell lineage in the fetal liver in the first trimester, but from the second trimester, the fetal bone marrow becomes the major site of B lymphopoiesis^7,8^. In contrast, in mice, the fetal liver remains the primary site of B lymphopoiesis until birth^9^. Further key differences exist between mice and human B lymphopoiesis including markers^10^, differentiation states^11^ (e.g., early lymphoid progenitors, ELPs and preProB), required cytokines (e.g., IL-7^12^), and specific signaling pathways (e.g., BTK, BLNK^13,14^). These species-specific distinctions underscore the need for a comprehensive characterisation of human B cell ontogeny, as well as the development of accurate human *in vitro* models.

A developmental block in B cell progenitor differentiation^15^, along with uncontrolled proliferation and increased susceptibility to chromosomal translocations that often promote interactions between B cell developmental genes and immunoglobulin heavy chain loci^16^, can predispose to leukemia^17^. Most paediatric B cell acute lymphoblastic leukemias (B-ALL) are thought to originate *in utero^18–22^*, with fetal ELPs identified as the cell of origin for the most common subtype of infant B-ALL (iB-ALL), driven by *KMT2A* rearrangements^23^. However, despite their central role in early detection, disease prevention, and therapeutic design, the fetal-specific gene programs that define early B lymphopoiesis, and those co-opted in paediatric leukemia, remain largely unexplored. Moreover, the TFs and regulatory networks that distinguish fetal from adult B lymphopoiesis, and potentially drive disease susceptibility, have yet to be systematically defined.

Here, we generated the most comprehensive single-cell multiomics atlas of human bone marrow hematopoiesis across prenatal and adult life, integrating single-cell RNA sequencing (scRNA-seq) and single-cell open chromatin (scATAC-seq) to reveal fundamental differences in cell states, chromatin dynamics, microenvironment interactions and leukemia-associated programs in B lymphopoiesis. Unlike prior stage-specific atlases^24^, our cross-lifespan dataset uncovers previously unrecognised features, including a novel ‘lateCLP’ population in adult bone marrow, reduced and highly similar V(D)J diversity in the prenatal stage, and distinct TF regulatory networks driving prenatal versus adult B lymphopoiesis. Importantly, fetal-specific gene programs and regulatory circuits identified in our atlas are reactivated in paediatric B cell leukemias, providing insights into the heightened susceptibility to malignant transformation of fetal-derived cells. Leveraging this atlas, we benchmarked a novel platform for generating B cells from human induced pluripotent stem cells (hiPSCs), which recapitulates *in vivo* developmental trajectories. This platform overcomes key limitations of prior OP9-based systems^25–28^, which we show fail to produce definitive HSCs, and establishes a scalable B cell production for disease modelling and future immunotherapies.

## Results

### A single-cell multiomics atlas of prenatal and adult bone marrow in humans identifies a novel lateCLP in adult bone marrow

Following the second trimester of fetal development, the bone marrow becomes a specialised niche for early B cell differentiation, supporting the progression of HSCs into progenitors that progressively develop into precursor B cells, which undergo immunoglobulin gene rearrangement and finally surface expression of the B cell receptor (BCR). To characterise this process, we generated a comprehensive single-cell multiomics atlas (scRNA-seq and scATAC-seq) by profiling 11 second-trimester fetal bone marrow samples (16-21 post conceptional weeks, pcw), when the bone marrow is the main hematopoietic organ producing B cells, and 6 adult bone marrow aspirates (38-82 years old, yo) (**Figure 1A, Supplementary Figure 1A-B**). ScATAC-seq was performed on 4 fetal and 4 adult bone marrow samples (**Figure 1B**).

**Figure 1.**
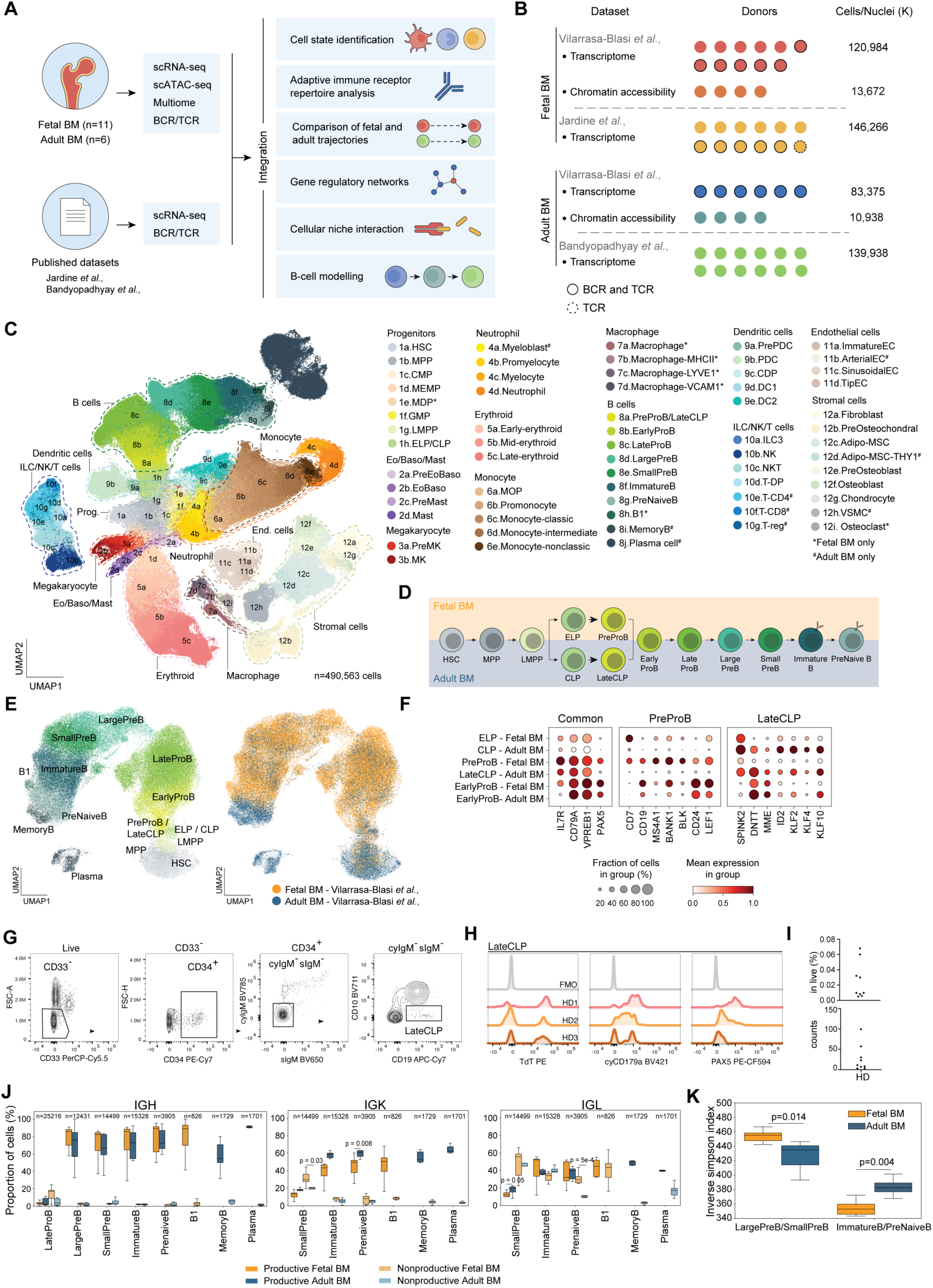
Single-cell multiomics atlas and B lymphopoiesis of the human fetal and adult bone marrow. A, Schematic of study design and analysis pipeline. scRNA-seq, scVDJ-seq, and scATAC-seq data were generated from fetal and adult bone marrow and integrated with publicly available scRNA-seq datasets^24,29^. The combined scRNA-seq and scATAC-seq atlas enabled: (1) characterisation of B cell states, (2) analysis of antigen receptor repertoire, (3) trajectory comparisons between fetal and adult B lymphopoiesis (4) construction of gene regulatory networks, (5) mapping of ligand-receptor interactions between B-cell and bone marrow niche, and (6) development of protocols for *in vitro* B cell modelling. **B,** List of datasets analysed, with donor and cell counts per sample. **C,** Batch-corrected Uniform Manifold Approximation and Projection (UMAP) of the integrated bone marrow scRNA-seq data from a total of 41 donors and 490,563 cells coloured by cell state (left) and developmental stage (right). **D,** Schematic illustration of the B-cell differentiation process in the fetal and adult bone marrow. Early lymphoid progenitors (ELPs) and preProB are only defined in the fetal bone marrow. Common lymphoid progenitors (CLPs) are only defined in the adult bone marrow. **E,** Batch-corrected Uniform Manifold Approximation and Projection (UMAP) embedding of the HSC differentiation to B cells containing exclusively cells sequenced for this study coloured by cell type (left) and their origin (right). **F,** Dot plot showing log-transformed, min-max normalised expression of marker genes for adult lateCLP cells. Differentially expressed between fetal and adult bone marrow: log-fold change >1, p-adjusted <0.05. **G,** Flow cytometric analysis of the lateCLP population, shown for one representative healthy adult bone marrow donor (n=9). **H,** Protein expression of TdT, cytosolic CD179a (cyCD179a) and PAX5 in lateCLP (gated according to panel ‘G’) of three healthy adult bone marrow donors (HD) compared to fluorescence minus one (FMO) control. **I,** Percentage of lateCLP cells in live cells (top) and absolute counts (bottom) of nine healthy adult bone marrow donors. **J,** Boxplot of the proportion of cells with productive or nonproductive heavy chain (IGH) and light chain (IGK or IGL). Orange colour represents fetal samples, blue colour represents adult samples. N, indicated number of cells. **K,** Box plot representing inverse simpson index for a group of early (largePreB and smallPreB) and late (immatureB and preNaiveB) B cell progenitors. Orange colour represents fetal samples, blue colour represents adult samples. BM, bone marrow; scRNA-seq, single cell RNA sequencing; sc-ATAC-seq, single cell sequencing assay for transposase-accessible chromatin; BCR, B cell receptor; TCR, T cell receptor; HSC, hematopoietic stem cell; MPP, multipotent progenitor; CMP, common myeloid progenitor; MEMP, megakaryocyte-erythorid-mast progenitor; MDP, monocyte-dendritic progenitor; GMP, granulocyte-monocyte progenitor: LMPP, lympho-myeloid primed progenitor; ELP, early lymphoid progenitor; CLP, common lymphoid progenitor; Eo, eosinophil; Baso, basophil; MK, megakaryocyte; MOP, myeloid progenitor; PDC, plasmacytoid dendritic cell; CDP, common dendritic cell; DC1, dendritic cell type 1; DC2, dendritic cell type 2; ILC3, innate lymphoid cell type 3; NK, natural killer; NKT, natural killer-T; T-DP, T cell double positive; T-reg, regulatory T cell; EC/Endo, endothelial; MSC, mesenchymal stromal cell; VSMC, vascular smooth muscle cell; Pre, precursor; Ig, immunoglobulin; IGH, immunoglobulin heavy locus; IGK, immunoglobulin kappa locus; IGL, immunoglobulin lambda locus.

After QC filtering, we retained 120,984 fetal and 83,375 adult cells from scRNA-seq with V(D)J data, as well as 13,672 fetal and 10,938 adult nuclei from scATAC. We further integrated our transcriptomics data with publicly available scRNA-seq data from the human fetal bone marrow^24^ (12-19 pcw) and adult bone marrow^29^ (52-74 yo), encompassing both immune and stromal compartments. This resulted in a unified dataset of 490,563 high quality cells - 267,250 fetal and 223,313 adult bone marrow (**Figure 1B-C, Supplementary Figure 1C-E, Supplementary Table 1-2**). Cell types in scRNAseq datasets were annotated based on the expression of well-characterised lineage markers and we used label transfer to annotate scATAC-seq data (**Supplementary Figure 2, Supplementary Figure 3A-B, Supplementary Table 3,** see **Supplementary Note 1-2**). We applied label transfer to compare our data with an additional inDrop-based scRNA-seq dataset comprising 28,702 cells from 5 paediatric (2-17 yo) and 10 adult (25-77 yo) bone marrow donors^30^, creating an independent dataset generated using a different technology to validate key findings (**Supplementary Figure 4A**).

To directly compare B cell differentiation processes between fetal and adult bone marrow while minimising batch effects, we performed dimensionality reduction and clustering analysis on cells spanning hematopoietic precursors to B cells (Vilarrasa-Blasi dataset) (**Figure 1D-E**). Plasma and memory B cells were identified in adults but absent in fetal bone marrow. This is consistent with previous reports showing the appearance of serum IgM only from 24 pcw^31^, suggesting that plasma cell development occurs later in gestation or in other anatomical sites. We identified a population of innate-like B1 cells exclusively in fetal bone marrow (**Supplementary Figure 1E**). While B1 cells have been previously identified in various fetal organs^32^, our single-cell transcriptomic analysis now reveals, for the first time, a substantial population of B1 cells in fetal bone marrow, characterised by the expression of canonical markers including *CD19*, *CD20*, *CD5*, *MZB1^33^* and *CCR10^32^*(**Supplementary Figure 2-4B**).

To compare transcriptomic differences between fetal and adult B cell development, we reconstructed differentiation trajectories from HSC to preNaiveB cells (also known as transitional cells) in both fetal and adult samples (**Supplementary Figure 4B-E, Supplementary Table 4, Supplementary Note 3**). The overall progression was similar in both cases: HSCs transition to multipotent progenitors (MPPs), which then give rise to lympho-myeloid primed progenitors (LMPPs). LMPPs commit to either early lymphoid progenitors (ELPs) predominant in the fetal bone marrow or common lymphoid progenitors (CLPs) predominant in the adult bone marrow. ELPs and CLPs clustered together (**Figure 1E**), but showed differential expression of defined markers. For example, *IL7R (CD127)* and *CD7^34,35^* were upregulated in ELPs, while *MME (CD10)^11^* was exclusively expressed in CLPs. Notably, *MME (CD10)* extended to later developmental stages (earlyProB up to immatureB) in both fetal and adult samples (**Figure 1F, Supplementary Figure 4B**).

In fetal bone marrow, differentiation from ELP to proB occurs predominantly via preProB cells (**Supplementary Figure 4E**), a population described as a hallmark of fetal B lymphopoiesis^11^. In adults, we defined a rare subset of adult cells clustering with preProB (**Figure 1E, Supplementary Figure 4F**) and positioned between ELP/CLP and proB in our trajectories (**Supplementary Figure 4E**) that we named ‘lateCLP’. LateCLP shared the expression of preProB markers *IL7R*, *CD79A* and *VPREB1^10,36^*. Compared to preProB, lateCLP presented lower expression of specific markers related to cell signaling such as *MS4A1 (CD20)*, *BANK1^37^*, *BLK^38^* and *CD24^39^*, as well as *CD7*, which is expressed at the ELP stage^11^ and correlates with the developmental decision between T-B and NK cell fate^40^, and *LEF1,* known to control precursor proliferation^41^. Conversely, lateCLP showed enhanced expression of *ID2* that regulates activity of E2A and was shown to increase in adult age^42^, as well as of genes related to control of proliferation and myeloid differentiation such as *KLF2*, *KLF4*, *KLF10^43–45^*, and of a hematopoietic progenitor gene as *SPINK2^46^*. LateCLP also expressed *MME (CD10)* and *DNTT (TdT)* at levels similar to CLPs, but lower than earlyProB cells **(Figure 1F, Supplementary Figure 4G).** We also demonstrated the presence of lateCLP, but not preProB cells, in paediatric scRNA-seq bone marrow samples, consistent with preProB cells being restricted to fetal development **(Supplementary Figure 4H)**.

To validate lateCLP as a novel cell population in the postnatal bone marrow, we developed a custom antibody panel and identified a population in the adult bone marrow that expressed CD34 (HSC marker maintained in early B cell differentiation stages), CD19 and TdT, with low levels of CD10, and the absence of immunoglobulin expression (**Figure 1G-I, Supplementary Figure 4I-K**). Adult lateCLP showed an intermediate phenotype between CLP and proB cells (**Supplementary Figure 4I-K**). The transcriptomic data revealed lower RNA expression of the canonical B cell marker CD19 in lateCLP compared to preProB (**Figure 1F**). The detectable presence of CD19 protein in lateCLP is supported by PAX5 expression, a TF crucial for B cell commitment and CD19 activation^47^.

Hence, our comprehensive map of B lymphopoiesis across prenatal and adult samples enabled the discovery of lateCLP cells as an adult-specific intermediate between CLP and proB cells. While sharing features with fetal preProB cells, lateCLPs exhibit a distinct transcriptional profile, making a previously unrecognised stage in human B cell development.

### Developmental divergence in V(D)J recombination reveals a stereotyped, public fetal B cell repertoire

The final phase of our inferred B cell trajectory involved the transition of lateProB cells to preB (large and small) followed by immatureB cells and preNaiveB cells (**Supplementary Figure 4E**). The transition from lateProB to preB was marked by the expression of the Ig heavy chain (IgH), part of the V(D)J recombination process. We reconstructed V(D)J recombination using Dandelion^48^ and included B1 cells (present only in fetal bone marrow), memoryB cells, and plasma cells (exclusive to adult bone marrow), since all these populations undergo BCR recombination.

Our analysis of V(D)J rearrangements showed the expected progression of immunoglobulin gene rearrangements through the different developmental stages starting in lateProB cells (**Supplementary Figure 5A**). The IgM heavy chain was productively rearranged from largePreB cells onwards in both fetal and adult bone marrow. We detected more IgD transcripts in preNaiveB cells in adults and the remaining isotypes (IgA and IgG) were exclusively present in antigen-experienced memoryB and plasma cells in adult bone marrow (**Supplementary Figure 5A)**. As expected^49,50^, we observed the presence of productive IGH rearrangements from the largePreB stage and light-chain IGK and IGL from smallPreB cells. The higher frequency of nonproductive IGK rearrangements in fetal smallPreB cells and persistence of nonproductive IGL rearrangements in preNaiveB cells in the fetal bone marrow suggested ongoing receptor editing at these stages **(Figure 1J)**.

In fetal bone marrow, IGH chain diversity was higher at earlier stages (largePreB / smallPreB cells), while in adult bone marrow, greater diversity was observed in the productive repertoire (immatureB / preNaiveB cells) (**Figure 1K**). In line with previous reports^51^, we observed a higher rate of N-nucleotide incorporation in the DJ junction in adults. In contrast to previous bulk analyses^51^, we observed no significant difference in the NP-nucleotide length in the VD junction between fetal and adults (**Supplementary Figure 5B)**. This discrepancy was likely due to the higher number of cells we analysed and the specific analysis of V(D)J recombination events in distinct B cell precursor populations. The total junction length of IGH was longer in adult cells **(Supplementary Figure 5B).** Longer CDR3 (complementarity determining region) **(Supplementary Figure 5B)** results from the addition of NP nucleotides due to increased TdT activity at the D-J junction in adults, as well as preferential use of proximal D-segments, that are shorter in sequence, in fetal B cells.

We observed a higher overlap of repertoires in fetal samples, as shown by the higher Morisita index among the individual samples (**Supplementary Figure 5C-D**). This reflects greater similarity in BCR repertories from fetal samples, which may be due to lower N-nucleotide incorporation and the preferential usage of conserved IGH genes. The more primitive fetal repertoire was characterised by the predominant usage of the D_H_-proximal IGHV1-2 and IGHV6-1, as well as the IGHD7-27 (**Supplementary Figure 5E**) as previously described^51–53^. While shared IGH clonotypes have been previously reported among different fetal samples^54^, here we observed similarity across individuals on a larger dataset that suggested a formation of ‘public’ or ‘convergent’ repertoire, largely overlapping among different individuals in the fetal stage. Functionally, this shared fetal BCR repertoire, characterised by reduced diversity and the use of conserved IGH genes, has been reported to bind apoptotic debris and commensal bacteria. It is also polyreactive but lacks the tolerance checkpoint between preNaive and naive B cells^55^, suggesting that peripheral B cell tolerance mechanisms are not yet fully developed at this stage.

Together, our analysis reveals fundamental developmental differences in V(D)J recombination between fetal and adult B cell precursors, including distinct timing of receptor editing, isotype expression, and junctional diversity. Notably, we demonstrate that the fetal B cell repertoire is less diverse but more stereotyped and convergent across individuals, forming a ‘public’ repertoire with potential innate-like functions.

### Fetal B lymphopoiesis transcriptional and microenvironmental programs are co-opted in leukemias

B lymphopoiesis is a multistep process orchestrated by TFs and modulated by extracellular signals from the surrounding microenvironment^56^. To identify TFs dynamically activated along fetal and adult B lymphopoiesis, we retrieved chromatin-accessible regions (peaks) specific to the B cell lineage for prenatal and adult samples, and combined them to reconstruct enhancer-mediated gene regulatory networks (GRNs) with SCENIC+^57^ (**Supplementary Figure 3B, Supplementary Figure 6A and 8A**, see **Methods**). Additionally, we linked these regulatory differences to signals from the microenvironment by identifying ligand-receptor interactions using CellPhoneDB v5^58^ (see **Methods, Supplementary Table 5**).

We began by examining the TFs activated during fetal B lymphopoiesis. As expected from the shared developmental trajectories of fetal and adult B cell differentiation, we identified canonical regulators of B cell differentiation, PAX5, TCF3 (E2A), FOXO1 and IKZF3 (Aiolos), in both contexts (**Supplementary Figure 6A and 8A**). However, fetal samples showed differential upregulation and activity of TFs linked to leukemia, including IKZF2^59^, TCF4^60^, EBF1^61^, LEF1^62^ and ZEB2^63^, particularly during early B cell developmental stages, from ELP to preProB cells (**Figure 2A-B, Supplementary Figure 6B, Supplementary Table 6**); all findings were are also found using inDrop-based paediatric and adult bone-marrow datasets (**Supplementary Figure 6C**). Interestingly, TFs upregulated during fetal B cell development were also linked to genetic risk variants for hematological disorders, suggesting that disease-associated GRNs may revert to a fetal-like state during pathogenesis (**Supplementary Figure 6D**).

**Figure 2.**
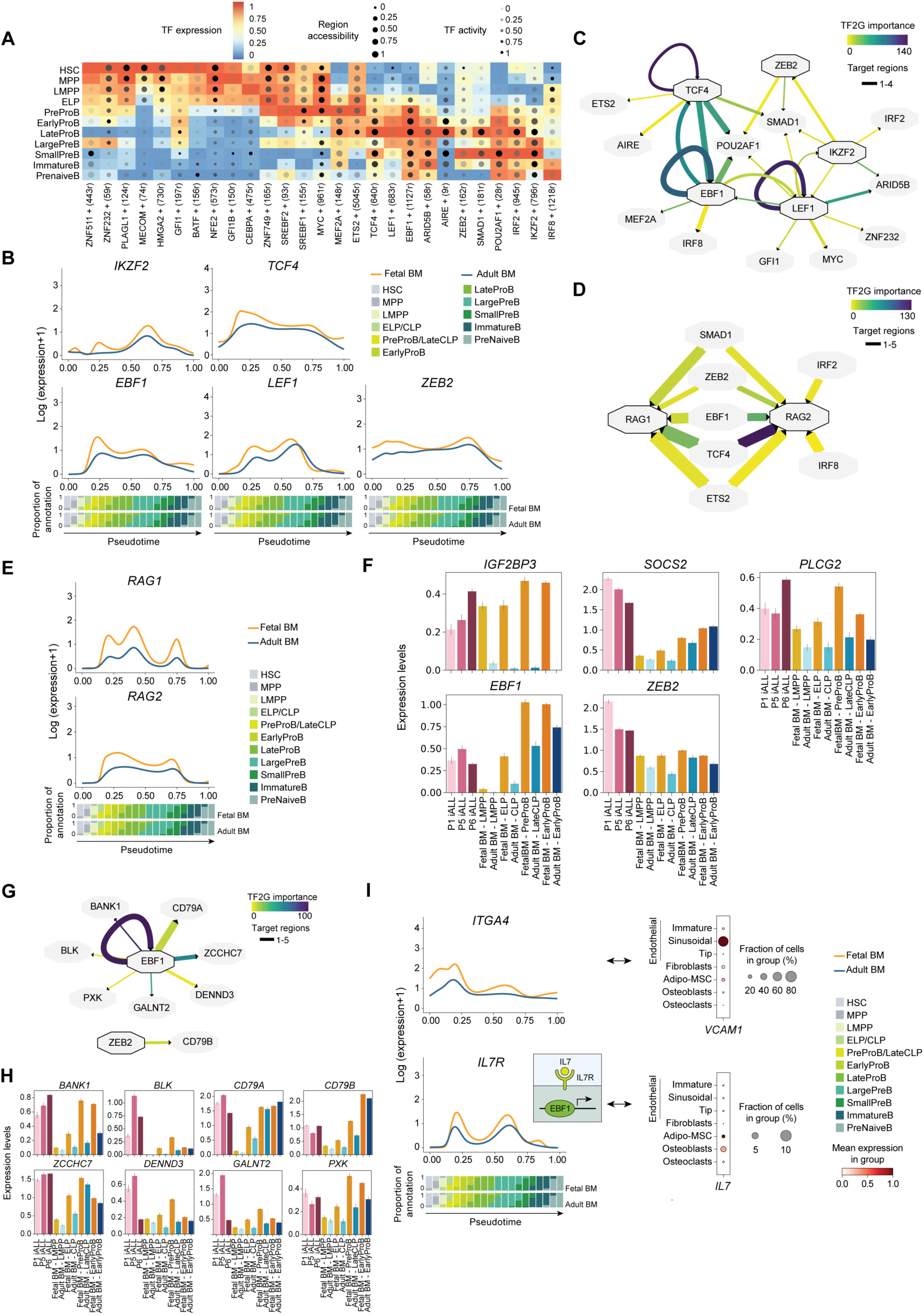
Regulatory programs upregulated in fetal B lymphopoiesis. A, Heatmap showing transcription factor (TF) activity during fetal B lymphopoiesis, based on scRNA-seq and scATAC-seq data (SCENIC+). Cell colour indicates TF expression, dot size reflects chromatin accessibility of TF eRegulon regions, and dot transparency represents overall TF regulon activity. Cell types are ordered by B cell differentiation. TFs shown are upregulated and differentially active in fetal vs. adult cell types. **B,** Smoothed spline curves showing gene expression dynamics of key TFs upregulated in fetal B lymphopoiesis. **C,** Subset of the SCENIC+ inferred gene regulatory network highlighting the regulatory relationships between the TFs shown in ‘A’ and ‘B’. **D,** Subset of the SCENIC+ inferred gene regulatory network showing the TFs that regulate the B cell recombination genes *RAG1* and *RAG2*, which are overexpressed in the fetal bone marrow. **E,** Smoothed spline curves showing expression dynamics of B cell recombination genes *RAG1, RAG2, IL7R* and *ITGA4A* upregulated in fetal B lymphopoiesis. **F,** Subset of the SCENIC+ inferred gene regulatory network showing the targets of EBF1 and ZEB2 that are differentially expressed between iB-ALL with KMTA-MLL1, KMTA-MLL2 and KMTA-MLL4 rearrangement compared to adult bone marrow. **G,** Bar plot showing the expression levels of the targets of EBF1 and ZEB2 upregulated in iB-ALL. **H,** Dot plot showing log-transformed, min-max normalised expression of two ligands (*IL7* and *VCAM1*) expressed in the stromal compartment of the fetal bone marrow binding IL7R and ITGA4A. Both interactions were identified with CellPhoneDB. **I,** Bar plot showing the expression levels of genes involved in proliferation and B cell differentiation that meet two criteria: (1) are upregulated in iB-ALL with KMTA-MLL1, KMTA-MLL2 and KMTA-MLL4 rearrangements when compared to adult CLPs (log-fold change >1, p-adjusted <0.01) and (2) are upregulated in fetal ELP when compared to adult CLP (log-fold change >0, p-adjusted <0.05) (related to Supplementary Figure 7). BM, bone marrow; HSC, hematopoietic stem cell; MPP, multipotent progenitor; CMP, common myeloid progenitor; LMPP, lympho-myeloid primed progenitor; ELP, early lymphoid progenitor; CLP, common lymphoid progenitor.

Regulons analysis indicated that EBF1, ZEB2 and TCF4 may influence the expression of *RAG1* and *RAG2*, genes critical for V(D)J recombination and implicated in leukemogenesis^64^ (**Figure 2D**). Transcriptomic data showed higher *RAG1* and *RAG2* expression along the fetal B cell trajectory compared to adults, with these findings also found using the inDrop-based paediatric and adult datasets (**Supplementary Figure 6C**). Interestingly, expression dynamics differed between developmental stages: in fetal bone marrow, *RAG2* expression peaked earlier, starting at the ELP stage, whereas in adults, the peak occurred later, at the lateCLP stage (**Figure 2E**).

To explore the potential link between TFs upregulated in fetal samples and leukemic gene expression programs, we reanalysed paediatric KMT2A-rearranged iB-ALL, where rearrangement events occur very early in development^23^. Using stringent filtering to minimise false positives, we found that *EBF1* and *ZEB2* were also upregulated in iB-ALL, along with proliferation-associated genes such as *IGF2BP3^65^* and *SOCS2^66^* (**Figure 2F**). Analysis of the EBF1 regulon revealed genes co-upregulated in fetal samples and iB-ALL, including genes relevant for B cell differentiation such as *BANK1, CD79A, BLK, ZCCHC7, PXK, DENND3* and *GALNT2.* Similarly, ZEB2 was inferred to regulate expression of the B cell co-receptor *CD79B* (**Figure 2G-H**). This suggests that sustained upregulation of *EBF1* and *ZEB2* in iB-ALL may contribute to the more aggressive phenotype of leukemias, reflecting the roles of these genes in fetal B lymphopoiesis.

Finally, we examined the influence of stromal interactions on GRNs throughout B cell differentiation. *ITGA4-VCAM1* and *IL7-IL7R* interactions were specifically upregulated during fetal B lymphopoiesis (**Figure 2I, Supplementary Table 5**). Receptors *ITGA4* and receptors were significantly upregulated in the fetal compared to adult B lymphopoiesis (**Figure 2I**), a finding validated using the inDrop-based paediatric and adult bone marrow dataset (**Supplementary Figure 6C**). Their cognate receptors were expressed by sinusoidal endothelial cells (*VCAM1*) as well as osteoblasts and adipo-MSC (*IL7*) (**Figure 2I)**. Notably, IL-7 positively regulates EBF1^35^ which shows enhanced activity during fetal B lymphopoiesis **(Figure 2A-B)**, further highlighting the unique regulatory environment of fetal B cell differentiation. Both interactions are critical for B cell commitment^67^, survival^35,68^ and proliferation of early precursors, and promote B cell fate by activating key regulators such as BACH2, PAX5 and EBF1^35^, all of which have been implicated in leukemia^69,70^.

Together, our findings reveal that fetal B lymphopoiesis is characterised by specific transcriptional and microenvironmental programs, such as sustained IL-7 signaling, early RAG activity, and the activation of EBF1 and ZEB2 regulons, that are also active in paediatric leukemia.

### Adult bone marrow rewires transcriptional networks to support B cell maturation and plasma cell niches

Using the same strategy on adult samples, we identified key TFs driving adult B cell differentiation (**Figure 3A**, **Supplementary Figure 8A)**. Several TFs regulating core cellular processes were upregulated during adult B lymphopoiesis (**Supplementary Figure 8B-D**). We previously defined *DNTT (TdT)* as a gene upregulated in the newly defined lateCLP state and maintained at high levels throughout B cell differentiation **(Figure 1G**, **Figure 3B)**. Here, we found that several TFs active in the lateCLP state, including MAZ, TCF4, FOXO1, E2F2, SMAD1, MXD3, NR3C1, target *DNTT* **(Figure 3C)**. Additionally, E2F2, FOXO1, IRF4, MEIS1, MXD3, NR3C1, SMAD1, TCF3 and TCF4 targeted *MME (CD10)*, which was upregulated in CLP/lateCLP in respect to ELP/PreProB **(Supplementary Figure 8E)**.

**Figure 3.**
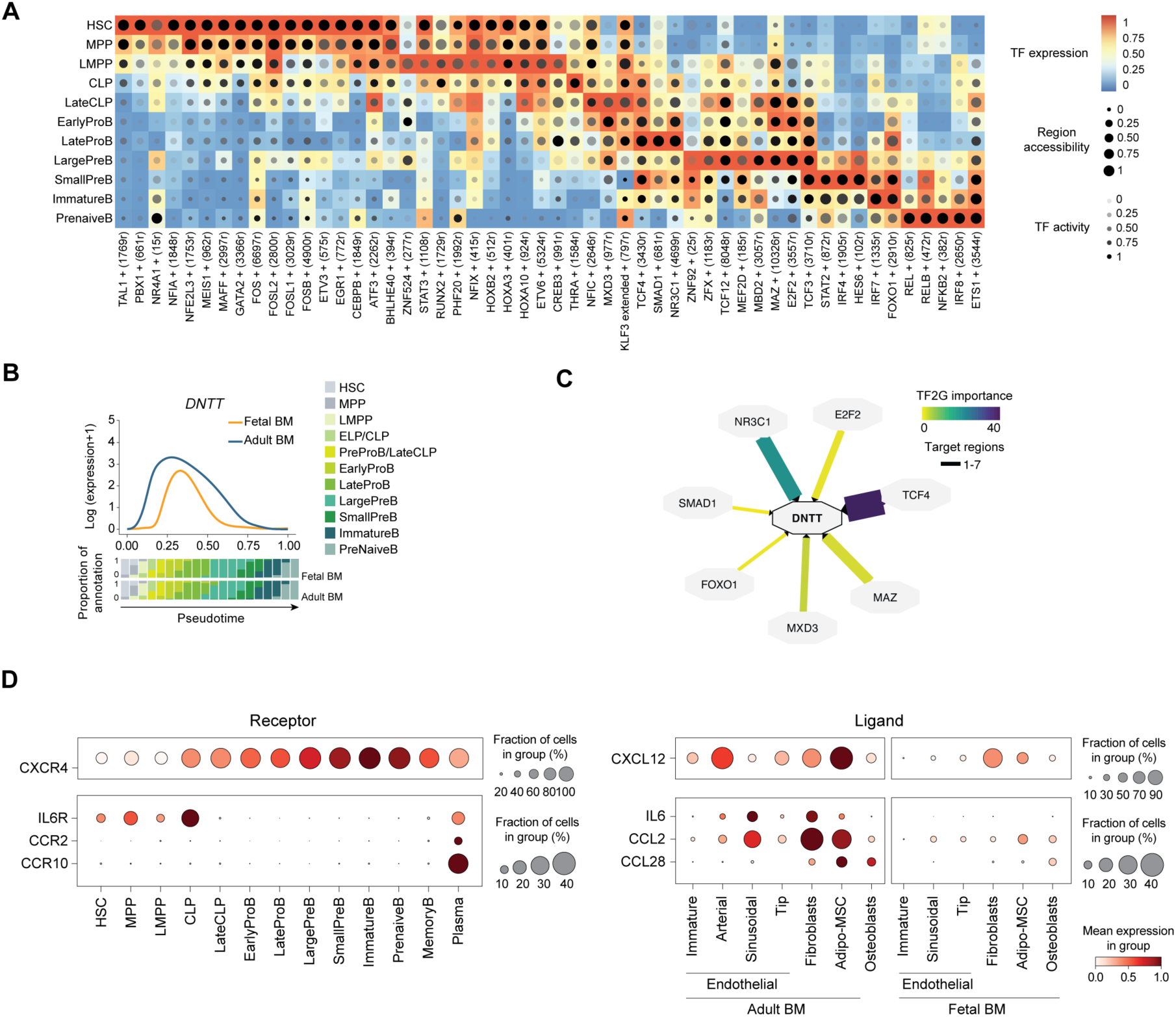
Regulatory programs upregulated in adult B lymphopoiesis. A, Heatmap showing transcription factor (TF) activity during adult B lymphopoiesis, based on scRNA-seq and scATAC-seq data (SCENIC+). Cell colour indicates TF expression, dot size reflects chromatin accessibility of TF eRegulon regions, and dot transparency represents overall TF regulon activity. Cell types are ordered by B cell differentiation. TFs shown are upregulated and differentially active in adults vs. fetal cell types. **B,** Subset of the SCENIC+ inferred gene regulatory network showing the TFs (from panel ‘A’) regulating DNTT involved in B cell V(D)J recombination. **C,** Smoothed spline curves showing *DNTT* gene expression dynamics during fetal and adult B lymphopoiesis. **D,** Dot plot showing log-transformed, min-max normalised expression of genes encoding cell-cell interaction molecules (y-axis) involved in B and plasma cell retention and function. Expression is shown across distinct B cell populations in adults (x-axis, left) and stroma/endothelium (x-axis, right). All genes are upregulated in adult samples. Cell-cell interactions were identified with CellPhoneDB. BM, bone marrow; HSC, hematopoietic stem cell; MPP, multipotent progenitor; CMP, common myeloid progenitor; LMPP, lympho-myeloid primed progenitor; ELP, early lymphoid progenitor; CLP, common lymphoid progenitor.

Next, we quantified extrinsic regulators of B cell progenitors in adults and found that *CXCR4-CXCL12* interaction between B cell progenitors (expressing *CXCR4*) and the stroma (adipo-MSC expressing *CXCL12*) was significantly upregulated in adult bone marrow (**Figure 3D)**. This interaction supports HSC survival, maintenance^71^, initial differentiation into B cells and retention of both HSCs and immatureB cells. In immatureB cells, the CXCR4/CXCL12 axis, together with PI3K signaling, regulates central tolerance by ensuring the selection of functional B cells^6,72^. Exit into circulation is mediated by *CXCR4* downregulation and *S1P* receptor upregulation^73^, both of which were higher in adult B cell progenitors compared to their fetal counterparts (**Supplementary Figure 8F-G**). Hence, in the adult microenvironment, retention signals contribute to the maintenance of tolerance and their reduction regulates B cell exit from the bone marrow and peripheral B cell replenishment.

Plasma cells are terminally differentiated B cells responsible for long term antibody secreting memory. While they are absent in fetal development up to 21 pcw (**Supplementary Figure 1E**), they are supported in adults by a specialized niche: cell-cell communication analysis revealed adult-specific stromal expression of *IL6^74,75^, CCL2^76,77^* and *CCL28^78,79^* (**Figure 3D**), cytokines and chemokines known to promote plasma cell survival. These signals were predominantly produced by sinusoidal cells, fibroblasts and adipo-MSC, with the corresponding receptors, *CCR2* and *CCR10*, expressed exclusively on plasma cells (**Figure 3D**).

In summary, our findings highlight that adult B lymphopoiesis is shaped by distinct transcriptional and microenvironmental programs that promote B cell retention, central tolerance, and the establishment of a plasma cell niche, which are absent during fetal development.

### Benchmarking hiPSC protocols established a framework for *in vitro* generation of B cell progenitors

Differentiation of hiPSCs offers a scalable model to study early B cell development and generate mature B cells *in vitro*. However, existing hiPSC-based protocols^25–28^ have shown low efficiency, and it remains unclear whether they recapitulate fetal or adult programs, or how feeder cells contribute to lineage outcomes. A comprehensive understanding of the *in vitro* modelling is essential to fully exploit its potential to study malignant transformation and drug targeting.

We benchmarked a published two-step protocol using OP9 and MS-5 stromal cells^28,80^, which the authors reported to yield CD19+ cells, although molecular and functional characterization of these cells was not provided. We followed the published protocol and generated scRNA-seq data on the output, we observed a strong myeloid bias and unreliable B cell differentiation, regardless of the hiPSC line used (**Supplementary Figure 9A-B**), consistent with findings from other groups^27^. To investigate whether this inefficiency was due to immature progenitors, we constructed an atlas of HSC maturation by integrating our *in vitro* data with *in vivo* datasets spanning key stages of fetal hematopoiesis: (1) human yolk sac^32,46,81–83^, where primitive HSC incapable of generating classical B cells emerges after ∼18 days post-conception; (2) aorta-gonad mesonephros (AGM)^46^, where definitive HSCs arise ∼ 4pcw; (3) fetal liver^24,32,46,81,83–85^, where HSCs expand after ∼5 PCW ; and (4) fetal bone-marrow (Vilarrasa-Blasi dataset and publicly available dataset^24^) (**Figure 4A-B, Supplementary Figure 9C**). Using this development-spanning data, we found that HSCs generated under OP9 conditions showed a primitive profile, marked by low expression of definitive markers including *CD34*, *SPINK2*, *ITGA4* and *CD74* (**Supplementary Figure 9D**).

**Figure 4.**
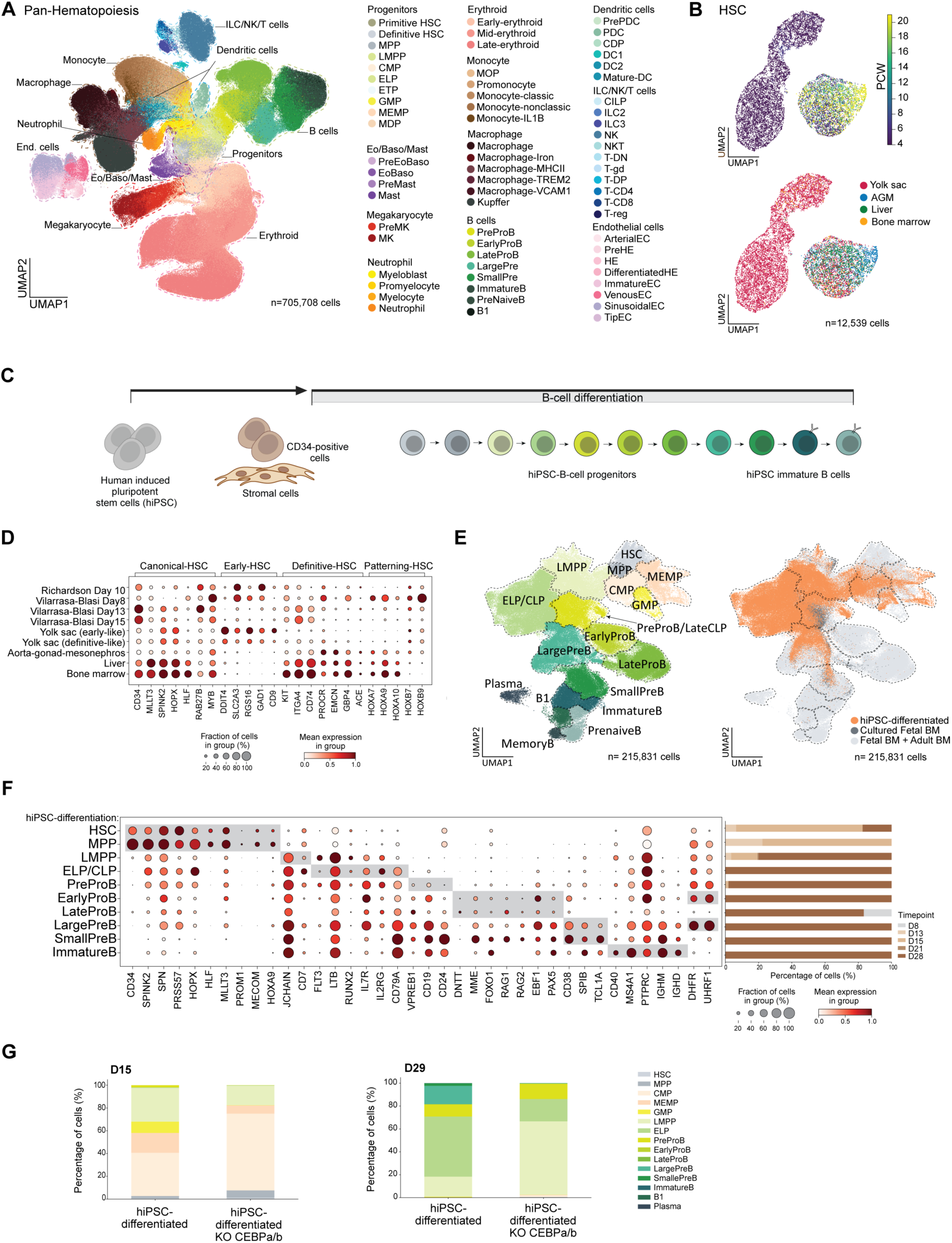
Characterisation of B cells differentiated from human induced pluripotent stem cells (hiPSC). A, Batch-corrected Uniform Manifold Approximation and Projection (UMAP) embedding of fetal hematopoiesis in both primitive hematopoiesis (yolk sac) and definitive hematopoiesis (aorta-gonad-mesonephros, liver and bone marrow) coloured by cell type (left). Total number of 705,708 cells. **B,** Batch-corrected UMAP embedding of hematopoietic stem cells (HSCs) coloured by post-conceptional weeks (pcw) (top) and organ (bottom). Total number of 12,539 cells. **C,** Schematic illustration of Vilarrasa-Blasi protocol to produce hiPSC-derived B cells. **B,** Batch-corrected Uniform Manifold Approximation and Projection (UMAP) embedding of fetal hematopoiesis in both primitive hematopoiesis (yolk sac) and definitive hematopoiesis (aorta-gonad-mesonephros, liver and bone marrow) coloured by cell type (left). Total number of 705,708 cells. **C,** Batch-corrected UMAP embedding of hematopoietic stem cells (HSCs) coloured by post-conceptional weeks (pcw) (top) and organ (bottom). Total number of 12,539 cells. **D,** Dot plot showing log-transformed, min-max normalised expression of primitive and definitive HSCs markers in both HSC derived from Richardson *et al.,* and our protocol. **E,** UMAP embedding of hiPSC-differentiated cells together with cultured fetal bone marrow cells and fetal and adult bone marrow. Coloured by cell lineage (left) and experimental model (hiPSC-differentiated: orange, cultured fetal bone marrow: dark grey tone and fetal and adult bone marrow: light grey tone). Total number of 215,831 cells. **F,** Dot plot showing log-transformed, min-max normalised expression of gene markers per cell type of hiPSC-derived B cells. Bar plot showing the percentage of cells for each cell state at each time point of the experiment (right). **G,** Bar plot showing percentage of cells in hiPSC-differentiated and hiPSC-differentiated CEBPA/B knockout experiments at time point day 15 and day 29. BM, bone marrow; AGM, aorta-gonad-mesonephros; HSC, hematopoietic stem cell; MPP, multipotent progenitor; MPP, lympho-myeloid primed progenitor; CMP, common myeloid progenitor; ELP, early lymphoid progenitor; ETP, early T cell; GMP, granulocyte-monocyte progenitor; MEMP, megakaryocyte-erythroid-mast progenitor; MDP, monocyte-dendritic progenitor; MOP, myeloid progenitor; PDC, plasmacytoid dendritic cell; CDP, common dendritic cell; DC1, dendritic cell type 1; DC2, dendritic cell type 2; DC, dendritic cell; CILP, common innate lymphoid cell; ILC2, innate lymphoid cell type 2; ILC3, innate lymphoid cell type 3; NK, natural killer; NKT, natural killer-T cell; T-DN, T cell double negative; T-gd, gamma delta T cell; T-DP, T cell double positive; Pre, precursor; EC/Endo, endothelial cell; HE, hemogenic endothelium; hiPSC, human induced pluripotent stem cell. The illustration from ‘Figure 5A’ was partially created using BioRender (https://biorender.com).

To overcome the limitations of OP9 based systems, we optimised a protocol and obtained CD19+ cells (**Figure 4C**). The resulting data showed that the *in vitro* derived HSCs had a definitive transcriptional profile, with high expression of *CD34*, *SPINK2*, *ITGA4* and *CD74* (**Figure 4D**).

We next characterised B cell precursors generated on mouse feeder cells by scRNA-seq, which were sorted by CD19 and CD10 at day 29, representing ∼1% of all CD45-positive immune cells (**Supplementary Figure 9F**). To our knowledge, this is the first molecular profiling of a B cell differentiation protocol from hiPSC. Our protocol generated all major stages of fetal B cell lymphopoiesis, including MPP (*CD34, SPINK2, SPN, PRSS57, HOPX, HLF, MLLT3, PROM1, MECOM, HOXA9*), LMPP (*JCHAIN, CD7, LTB, FLT3, LTB*), ELP (*RUNX2, IL7R, IL2RG, CD79A*), preProB (*VPREB1, CD19, CD24*), early and late proB (*DNTT, MME FOXO1, RAG1, RAG2, EBF1, PAX5*), large and small PreB (*CD19, MME, CD38, SPIB, TCL1A*) and *bona fide* immatureB cells (*CD40, MS4A1, PTPRC*) at low frequency (∼0.1% of all the immune cells) (**Figure 4E-F, Supplementary Table 8**). We also compared B cell output from hiPSCs and CD34+ fetal bone marrow cells, with and without depletion of CD19 and CD10 populations. Based on CD19 and CD10 expression at day 29, B cell differentiation efficiency ranged from 6-10% of immune cells in fetal BM cultures (**Supplementary Figure 9G-H**), and all key stages were similarly captured (**Supplementary Figure 9G-H**).

In addition to B cells, our hiPSC derived cultures contained various myeloid lineages, including monocytes, macrophages, dendritic cells and mast cells. To investigate whether myeloid and B cell differentiation pathways influence each other, we knocked out CEBPA and CEBPB, blocking myeloid development at the CMP stage. Despite reduced frequency, B-cell specification was maintained (**Figure 4G, Supplementary Figure 9I**), suggesting that myeloid priming does not compromise B cell potential.

This supports current models proposing progressive and flexible lineage specification^87^, consistent with findings that *CEBPA* overexpression can redirect B cell precursors to a myeloid fate^88^.

In sum, we have established a protocol for *in vitro* B cell differentiation from hiPSCs that recapitulates key stages of human B lymphopoiesis. This system enables the generation of definitive HSCs and B cell progenitors, offering a tractable model for studying human B cell development and disease.

### Stromal signals shape transcriptional dynamics during *in vitro* B lymphopoiesis

We next studied *in vitro* B cell differentiation dynamics, comparing our protocol with fetal and adult B lymphopoiesis. Using Slingshot for trajectory inference and aligning with the Genes2Genes^89^ framework, we mapped gene expression changes along matched developmental paths between *in vitro* and *in vivo* systems (**Figure 5A, Supplementary Figure 10A-B**).

**Figure 5.**
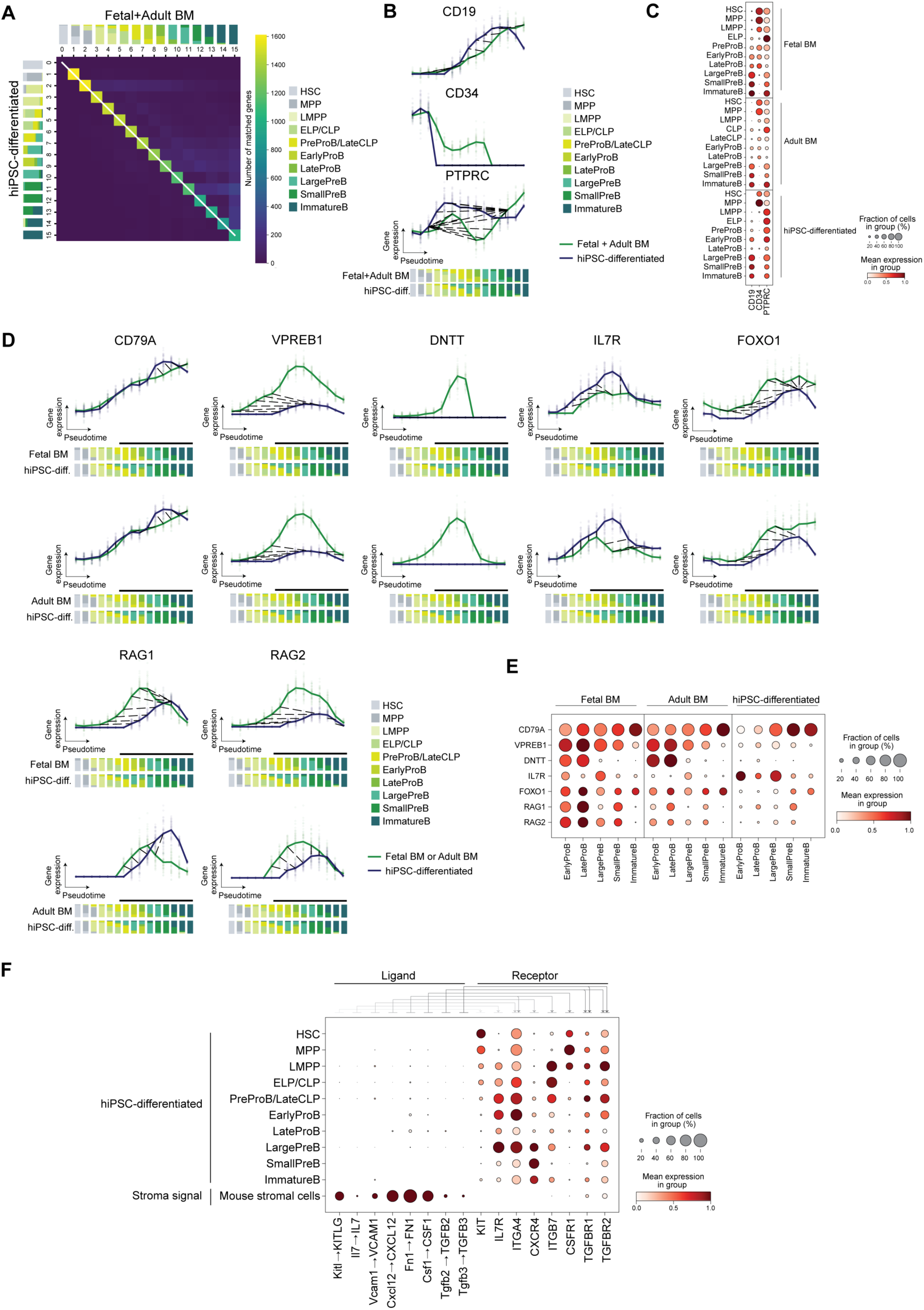
hiPSC derived B cells recapitulate fetal B lymphopoiesis. A, Pairwise time point matrix between the reference (fetal and adult BM) and the query (hiPSC-differentiated) of transcription factors and B related cell genes. Color represents total gene count showing a match between corresponding time points. White line represents the average alignment path. Stacked bar plots represent reference and query cell composition across 15 equispaced pseudotime points, colored by cell state. **B**, Gene expression of three markers (*CD19*, *CD34* and *PTPRC* (CD45) in hiPSC-differentiated (query, blue) and fetal and adult BM (reference, green). Interpolated log1p-normalized (per-cell total raw transcript counts normalized to 10,000 and log1p-transformed) expression (y-axis) against pseudotime (x-axis). Bold lines represent mean expression trends and faded data points indicate 50 random samples from the estimated expression distribution at each time point. Black dashed lines represent time point matches (captured by the alignment string below). Stacked bar plots represent reference and query cell composition across 15 equispaced pseudotime points, colored by cell state. **C,** Dot plot showing log-transformed, min-max normalised expression of markers shown in panel ‘B’. **D,** Gene expression of seven genes involved in rearrangement of the B cell receptor and immunoglobulin recombination (*CD79A*, *VPREB1*, *DNTT*, *IL7R*, *FOXO1, RAG1* and *RAG2)* in hiPSC-differentiated (query, blue) and fetal or adult BM (reference, green). Interpolated log1p-normalized (per-cell total raw transcript counts normalized to 10,000 and log1p-transformed) expression (y-axis) against pseudotime (x-axis). Bold lines represent mean expression trends and faded data points indicate 50 random samples from the estimated expression distribution at each time point. Black dashed lines represent time point matches (captured by the alignment string below). Stacked bar plots represent reference and query cell composition across 15 equispaced pseudotime points, colored by cell state. A black line above the stacked bar plots highlights the bars containing the cell states from earlyProB to immatureB. **E,** Dot plot showing log-transformed, min-max normalized expression of seven genes involved in the rearrangement of the B-cell receptor and immunoglobulin recombination, spanning from earlyProB to immatureB stages in fetal and adult bone marrow, as well as hiPSC-derived cells. **F,** Dot plot showing log-transformed, min-max normalised expression of ligands expressed by mouse stromal cells and receptors in hiPSC-B cell differentiated subpopulations. Arrows represent the ligand-receptor communications. BM, bone marrow; hiPSC, human induced pluripotent stem cell.

Canonical markers of B cell differentiation, including *CD34*, *PTPRC (CD45)* and *CD19*, were expressed in hiPSC-derived cells but with distinct expression dynamics (**Figure 5B-C**). *CD34* expression decreased sharply at the LMPP stage *in vitro*, while *in vivo* downregulation was more gradual, only nearing complete suppression at the largePreB stage. Indeed, the loss of *CD34* expression is a common observation in protocols to differentiate cells in culture^90,91^. In contrast, *PTPRC (CD45)* upregulation began earlier *in vitro*, around the LMPP stage, compared to its initiation *in vivo* at the ELP/CLP stage, and its expression levels remained higher *in vitro* throughout the process. *CD19* upregulation followed a comparable trend but was downregulated towards the end time points of the *in vitro* cultures. These differences suggest accelerated differentiation kinetics *in vitro*, likely due to the condensed 21-day culture timeline.

We also observed reduced expression of key immunoglobulin recombination genes, such as *DNTT (TdT), VPREB1*, *RAG1* and *RAG2* (**Figure 5D-E, Supplementary Figure 10C**). Low levels of *RAG2* expression were found at the lateProB and smallpreB cell stage, in line with lowered *FOXO1* expression that is crucial to induce *RAG^92^* **(Figure 5D-E, Supplementary Figure 10C**). Persistent IL-7 receptor signaling, as suggested by the increased and persistent *IL7R* expression *in vitro*, may contribute to the inefficient *FOXO1* induction^93^ (**Figure 5D-E, Supplementary Figure 10C**).

To understand the microenvironmental influence on these altered *in vitro* differentiation dynamics, we profiled signaling molecules expressed by mouse stromal cells (see **Methods**). Those stromal cells expressed low levels of IL-7, which was released into the media, and expressed crucial factors supporting more primitive progenitors such as *KITLG*, factors supporting proB and committed B cell precursors such as *CSF1*, and niche localization signals such as *VCAM1*, *CXCL12*, and *FN1* (**Figure 5F**). They also expressed *TGFbeta2* and *3*, established regulators of B lymphopoiesis which may contribute to the low output of immatureB cells in our culture^94^. Importantly, the cognate receptors for these molecules were expressed by the differentiating hiPSC-B cells. Unlike *in vivo* conditions, where lymphoid progenitors transition through distinct niches, mouse stromal cells provide overlapping maintenance (e.g. *KITLG*) and differentiation (e.g. *CXCL12, CSF1*) cues throughout the culture period. This mixed signaling environment may limit proper stage-specific progression of B cells.

Finally, we evaluated the system’s utility for disease modelling. Genes implicated in blood malignancies were expressed in our cultures (**Supplementary Figure 10D**), implying it can be a promising tool for disease modelling.

In summary, our findings show that mouse stromal cells provide essential signals that support early B cell development *in vitro* and can also alter transcriptional dynamics compared to *in vivo* differentiation, offering insights for future protocol refinement. Despite limited maturation, the system captures key precursor stages and expresses genes relevant to B cell malignancies, making it a valuable tool for studying human B lymphopoiesis and disease.

## Discussion

Understanding the regulation of human B lymphopoiesis is essential for addressing diseases ranging from leukemia to autoimmunity. Here, we present a high-resolution, multimodal atlas of fetal and adult bone marrow B cell development, revealing transcriptional, epigenetic, and niche-driven differences that shape immune function. Our findings reveal how fetal programs are co-opted in paediatric leukemia, highlighting potential drivers of disease aggressiveness. Alongside, we develop and validate a new protocol that captures definitive HSC emergence and all major stages of B cell differentiation *in vitro*. Together, these tools provide a valuable resource for dissecting the developmental origins of B cell function and transformation.

First, by comprehensively analysing all progenitor populations across prenatal and postnatal development, we identified lateCLP, a previously unrecognized population in adult bone marrow. This subset shares features with fetal preProB cells, including *IL7R*, *CD79A*, and *VPREB1* expression, but differs in reduced expression of early B cell markers like *CD20*, *BLK*, and *BANK1*. LateCLP cells expressed *CD10* and moderate levels of *PAX5* expression and protein supporting detectable *CD19* protein despite low transcript levels. Notably, these cells bridge the ELP/CLP and proB stages and are transcriptionally distinct from both, suggesting a functionally unique intermediate in adult B cell development.

Second, the enrichment of CD34+ progenitors in fetal datasets allowed us to confirm the presence of B1 B cells in fetal bone marrow. Although B1 cells have been previously described in human peripheral blood^95^ and other fetal organs, mainly in spleen^32^, their existence has been controversial^96,97^. Our single-cell data provide the first transcriptomic evidence for bona fide B1 cells in this compartment *in utero*, consistent with mice data^98^. These innate-like B cells, absent in adult marrow, likely contribute to the initial wave of IgM production in the fetus after 24 pcw^31^. As IgM does not cross the placenta, this endogenous antibody production provides both pre- and postnatal protection against pathogens.

Third, by reconstructing the immunoglobulin repertoire at single-cell resolution, we revealed developmental differences in V(D)J recombination and BCR diversity, extending previous findings from bulk or stage-specific analyses^51,53,99^. We confirmed the biased V(D)J usage in fetal B cells, shorter CDR3 length, and fewer N-nucleotide additions compared to adults, in line with lower *TdT (DNTT)* expression during fetal development^51–53^. Interestingly, prenatally, *DNTT* peaked at the lateProB stage rather than preProB, in contrast to mice, where TdT is absent during liver-based prenatal development, resulting in impaired junctional diversity^100^. The later onset of *DNTT* (at lateProB) and increased *RAG* expression in fetal cells may explain the higher frequency of heavy-chain rearrangements in fetal lateProB cells. These features contribute to the extensive overlap of fetal B cell repertoire across individuals, a level of convergence not previously reported^54^.

The repertoire overlap prenatally, especially in immatureB and preNaiveB cells, suggests the presence of ‘public’ fetal B cell repertoire with limited antigen specificity. In adults, similar ‘public’ clones arise only in mature B cell responses^101–103^, whereas in fetal samples, the overlap spans broader precursor stages. Higher repertoire similarity in preterm infants compared to term infants further supports the idea of early clonal expansion against shared antigens^104^. The convergence appears driven by preferential use of proximal gene segments, rather than peripheral expansion. Stage-specific analysis revealed a drop in heavy-chain diversity between fetal preB and preNaiveB cells, which is a checkpoint stage marked by proliferation upon successful rearrangement and pairing with the surrogate light chain^105^. Interestingly, fetal preB cells showed higher diversity than adult counterparts, possibly reflecting lower proliferation rates in fetal B cell development.

Fourth, we identified TFs and microenvironmental signals active in fetal B lymphopoiesis that are reactivated in paediatric leukemia. Notably, TFs such as EBF1 and ZEB2 upregulated in fetal progenitors were also elevated in KMT2A-rearranged iB-ALL. These TFs regulate key genes involved in BCR signaling (*CD79A/B*) and recombination (*RAG1, RAG2*). Increased RAG activity, coupled with high EBF1 and FOXO1 activity, has been associated with chromosomal translocations and leukemic transformation^16^. In addition, fetal-specific signals such as *IL7* and *ITGA4–VCAM1* interactions were enriched and likely contributed to enhanced EBF1 activity, a TF known to drive leukemogenic programs when dysregulated. These findings reinforce the concept of onco-fetal reprogramming^106^, where tumours reactivate fetal gene circuits that may promote aggressive disease phenotypes.

Fifth, we applied insights from our atlas to benchmark and improve protocols for in vitro B cell differentiation from hiPSC. Our benchmark revealed that existing OP9 and MS-5 based methods failed to generate definitive HSCs, consequently, robust B cell output, consistent with prior reports^27^. We optimised a protocol and successfully recapitulate all major stages of fetal B cell development, including MPP, ELP, preProB, proB, PreB, and immatureB cells. Molecular profiling showed accelerated yet stage-faithful progression, although key regulators like RAG2 and FOXO1 were downregulated, likely due to persistent IL-7 signaling and mixed feeder-derived cues. Importantly, this system also expressed genes implicated in leukemia, demonstrating its potential for disease modelling. Despite this process, the use of feeder cells remains a limitation, as those deliver overlapping maintenance and differentiation signals that may impair stage-specific maturation. Nonetheless, the development of a robust and well-characterised hiPSC-B cell protocol marks a significant milestone, providing a platform to dissect human B cell development, model lymphoid malignancies, and pave the way for the production of off-the-shelf antibody-producing cells.

In summary, we present two complementary resources: a multimodal atlas and an *in vitro* differentiation protocol that together reveal the developmental logic of human B lymphopoiesis, shed light on the fetal origins of paediatric leukemia, and enable future advances in disease modelling, immunotherapy, and regenerative medicine.

## Acknowledgements

This publication is part of the Human Cell Atlas– www.humancellatlas.org/publications/. We are thankful to the Sanger Core Sequencing pipeline for support with sequencing library preparation. We acknowledge Annie Moisan and Cara Buchanan, program director and coordinator from the Wellcome Leap HOPE program for their support. We thank Antonio García (https://www.bio-graphics.es/) for his invaluable help with conceptualising and making the illustrations that are part of this manuscript, and Aidan Maartens for proofreading and providing advice on the narrative of the manuscript. We extend our gratitude to Lisa Lindeboom for her assistance in setting up the project, and to Jessica Cox for her support with the CellPhoneDBViz portal. We also thank Bee Ling Ng and the Wellcome Sanger Institute Cytometry Core Facility for their support. The human embryonic and fetal material was provided by the Joint MRC/Wellcome Trust (grant no. MR/R006237/1) HDBR (http://www.hdbr.org). We thank the IR and FREEZE Biobank Freiburg at the University Medical Center Freiburg. The illustration from ‘Figure 5A’ was partially created using BioRender (https://biorender.com).

## Author information

R.V.-B. and R.V.-T. conceived and designed the experiments and analyses. R.V.-B. and S.K. analysed the data with contributions from P.P., K.T., V.L., H.A., M.P., L.G.-A. R.V.-B. and D.Z. performed sample processing with contributions from E.P. and C.S. R.V.-B. performed the hiPSC-B cell protocol with contributions from F.W., O.N., R.C., Y.S.M., I.K. R.V.-B., R.V.-T. and M.R. interpreted the data with contributions from J.K., M.S., L.G.-A, S.A.T., K.W., B.M.J, P.W.Z. R.V.-B., R.V.-T. and M.R. supervised the work and wrote the manuscript with contributions from L.G.-A. All authors read and approved the manuscript.

## Methods

### Patient samples

The human embryonic and fetal material was provided by the Joint MRC / Wellcome Trust (Grant #MR/006237/1) Human Developmental Biology Resource (http://www.hdbr.org) following elective of pregnancy, with written informed consent and approved by the London - Fulham Research Ethics Committee (REC reference 23/LO/0312). HDBR is regulated by the UK Human Tissue Authority (HTA; https://www.hta.gov.uk) and operates in accordance with the relevant HTA Codes of Practice. Adult bone marrow tissues were obtained from healthy donors that underwent hip replacement surgery and were otherwise healthy. All participants provided written informed consent in accordance with approval by the Ethics Committee of the University Medical Center Freiburg (20-1109).

### Tissue processing

Adherent material from the fetal bone marrow samples was removed, and the end of the bones were cut. Cells were recovered by flushing the bone marrow from one end. In a few samples, the bones were cut into pieces and ground with a pestle and mortar while adding buffer (PBS 5%FBS and 2mM EDTA) to reduce clumping^24^. For both collection methods, cells were filtered with a 70µm filter and centrifuge for 5 minutes at 500xg. After centrifugation, red blood cells were lysed by treating with RBC lysis buffer according to manufacturer’s instructions (eBioscience #00-4300). Finally, cells were counted and processed in one or more of the following way; (1) sequencing the whole population, (2) enriching for CD34-positive cells using the CD34-positive selection kit (Miltenyi Biotec #130-046-702), following manufacturer’s instructions, and sequencing, (3) enriching for CD34-positive cells and sorting for LMPP and MPP subpopulations^11^. Staining antibodies: Lin-FITC: CD2 clone: RPA-2.10 (BioLegend), CD3 clone OKT3 (BioLegend), CD14 clone M5E2 (BioLegend), CD16 clone 3G8 (BioLegend), CD56 clone HCD56 (BioLegend), CD235a clone Ga-R2/HiR2 (BD Biosciences); CD34-BV786 clone 581 (BD Biosciences); CD38-BV711 clone HIT2 (BioLegend); CD90-APC-Cy7 clone 5E10 (BioLegend); CD45RA-APC clone HI100 (BioLegend), and DAPI as live death dye. After sorting, the specific subpopulations were sequenced, (4) for one sample we enriched for CD45-positive cells using the CD45-positive selection kit (Miltenyi Biotec #130-045-801), following manufacturer’s instructions, and sequenced, (5) enriching for CD34-positive cells and depleting CD19-positive cells following published sorting strategy^11^. Staining antibodies: Lin-FITC: CD2 clone: RPA-2.10 (BioLegend), CD3 clone OKT3 (BioLegend), CD14 clone M5E2 (BioLegend), CD16 clone 3G8 (BioLegend), CD56 clone HCD56 (BioLegend), CD235a clone Ga-R2/HiR2 (BD Biosciences); CD34-BV786 clone 581 (BD Biosciences); CD19-PE clone HIB19 (BD Bioscience); CD10-APC clone HI10A (BD Biosciences)and DAPI as live death dye. After sorting, the CD19-negative population was cryopreserved, and/or (6) cryopreserving and storing at -80°C for 24 hours before transferring to liquid nitrogen.

Adult bone marrow samples were collected in PBS + 0.1% EDTA before being centrifuged for 10 minutes at 150xg to remove the remaining fat layer. The samples were then mashed and filtered using a 70µm filter. Mononuclear cells were isolated by density gradient centrifugation (20 minutes, 1000xg) and washed afterwards with PBS + 0.1% EDTA (7 minutes, 300xg). Cells were sorted to enrich our analysis for particular populations using Bigfoot spectral cell sorter (ThermoFisher scientific). Briefly, cells were incubated with antibodies listed in **Supplementary Table 9** together with Zombie NIR live/dead fluorescence dye (BioLegend #423106) in a total volume 150µl (PBS, 2% heat inactivated FBS, 1 mM EDTA) for 15 minutes on ice. After washing the cells, sample was sorted to obtain the CD90-positive and CD34-positive cells and enrich for CD45-positive CD33-negative cells (50%), CD33-positive cells (25%) and CD45-negative CD33-negative cells (25%). Data were analysed using FlowJo Software (v10.10.0).

### Flow cytometry of adult bone marrow samples

Phenotyping of cells was performed with antibodies listed in the **Supplementary Table 9**. Intracellular proteins were stained with eBioscience Foxp3/Transcription Factor Staining Buffer Set (Invitrogen) according to the manufacturer’s protocol. Briefly, cells were stained with antibodies for surface protein detection and after washing, 100µl of Foxp3 Fixation/Permeabilization working solution was added for 30 minutes and incubated at room temperature. After two washing steps with 1x Permeabilization Buffer, the intracellular proteins were stained for 30 minutes at room temperature. The cells were washed twice with 1x Permeabilization Buffer prior acquisition. Samples were acquired on Cytek Aurora (Cytek Biosciences) with SpectroFlo Software version 2.2.0 and 3.1.0. Data was analyzed by FlowJo Software (v10.7, TreeStar).

### Cell culture

iPS11 hiPSCs (ALSTEM Inc, kindly gifted by Zandstra’s lab) were cultured in mTeSR-plus media (STEMCELL Tech #100-0276) on 10cm plates (Corning #430167) coated with hESC-qualified Matrigel (Corning #354277) at a final concentration of 180µg/mL, diluted in 6mL DMEM/F12 (Gibco #21331020). Plates were coated overnight in the fridge, wrapped with parafilm, and brought to room temperature before use. iPS11 hiPSCs were passaged as clumps when 60-70% confluency is reached by washing with PBS (Gibco #20012027) before treating with 4mL of TrypLE Express (Gibco #12605-010) at 37°C for 5 minutes. TrypLE was gently aspirated and cells were scraped down into mTeSR-plus media using a cell scraper (Sarstedt #83.3951) and plates were washed with fresh mTeSR-plus media. Gently pipette mix 1-3 times to break up remnant large colonies and seed on freshly Matrigel-coated plates. Cells were maintained at 37°C, 5% CO_2_. Media was changed daily.

Kolf_2 (HipSci) were cultured in E8-flex (Life Techologies #A2858501) on 10 cm plates (Corning #430167) coated with Vitronectin XF matrix (VTN, STEMCELL Technologies #07180) at a 1:25 dilution with PBS (Gibco #10010015). Plates were coated for an hour at 37°C. Kolf_2 were passaged using ReLesR (STEMCELL Tech #100-0483) at 37°C for 3 minutes. An appropriate volume of media was added, and colonies were suspended to achieve a uniform size before seeding. Cells were maintained at 37°C, 5% CO_2_. Media was changed daily.

MIFF3 hiPSC (UOSi001-B, University of Sheffield) were culture in E8 (Life Technologies #A1517001) on 10cm plates (Corning #430167) coated with Vitonectin-N recombinant human protein (VTN-N, LifeTechnologies #A14700) in a dilution 1:100 with DPBS Ca^2+^/Mg^2+^ free (Gibco #14190094). Plates were coated for an hour at room temperature. MIFF3 cells were passaged using ReLeSR (STEMCELL Technologies #100-0483). Cells were washed with ReLeSR, the solution was then aspirated, and colonies were left for 5 minutes to detach. An appropriate volume of media was added, and colonies were suspended to achieve a uniform size before seeding. Cells were maintained at 37°C, 5% CO_2_. Media was changed daily.

A18966, gibco human episomal iPSC line (ThermoFisher) were cultured in E8 (Life Technologies #A1517001) on 10cm plates (Corning #430167), coated with Vitronectin XF matrix (VTN, STEMCELL Technologies #07180) in a dilution 1:50 with DPBS (Gibco #14190094). Plates were coated for an hour at 37°C. A18966 cells were passaged using ReLesR (STEMCELL Technologies #100-0483) at 37°C for 4-5 minutes. An appropriate volume of media was then added, and colonies were suspended and seeded to achieve a uniform colony size. Cells were maintained at 37°C, 5% CO_2_. Media was changed daily.

### Derivation of CEBPa and CEBPb double knock-out hiPSCs

Stock gRNAs for *CEBPa* and *CEBPb* (Synthego Gene Knockout Kit V2 1.5nmol comprising 3x equimolar gRNAs per gene) were prepared in 1X TE pH 7.5 solution (IDTE, IDT #11-01-02-02) at 200pmol/µL (200µM). 22.5pmol of each gRNA mix (a total of 45pmol) were assembled into ribonucleoproteins (RNPs) complexes with 20pmol of eSpCas9NLS (NEB #M0606T) and left at room temperature for 20 minutes inside the tissue culture flow hood. Cultured iPS11 iPSCs of early passage (p7) on a 10cm plate were isolated as single cells by washing with PBS and treating with 4mL of TrypLE Express (Gibco #12604013) at 37°C for 2.5 minutes. Gently aspirate TrypLE without dislodging colonies before resuspending cells and washing the plate once in a total of 10mL of mTeSR-plus media. Single-cell density was checked and counted by diluting equivolume with 0.4% trypan blue solution (Gibco #T10282) using a haemocytometer (VWR #631-1098). 200K cells were centrifuged at 300xg for 3 minutes and supernatant was aspirated. Cell pellet was resuspended in 16.4µL P3 buffer + 3.6µL P3 supplement + 2µL RNP mix and transferred to a 16-well strip cuvette (Lonza #V4XP-3032) for nucleofection using the CA-137 program in a 4D-Nucleofector-X unit (Lonza). Cells were immediately transferred to room temperature-equilibrated mTeSR-plus before seeding in a serial dilution on a Matrigel-coated 6-well plate in mTeSR-plus media supplemented with 10µM Y-27632 (Merck #Y0503-1MG). The cells were cultured for approximately 10 days to allow the formation of single-cell colonies. Colonies were picked by treating each well with 1mL of gentle cell dissociation reagent for 2-3 minutes at room temperature on a microscope until cell-cell adhesions were weakened without dislodging. Bent filtered P10 tips were used to pick colonies into a Matrigel-coated well of a 96-well plate containing mTeSR-plus supplemented with 10µM Y-27632.

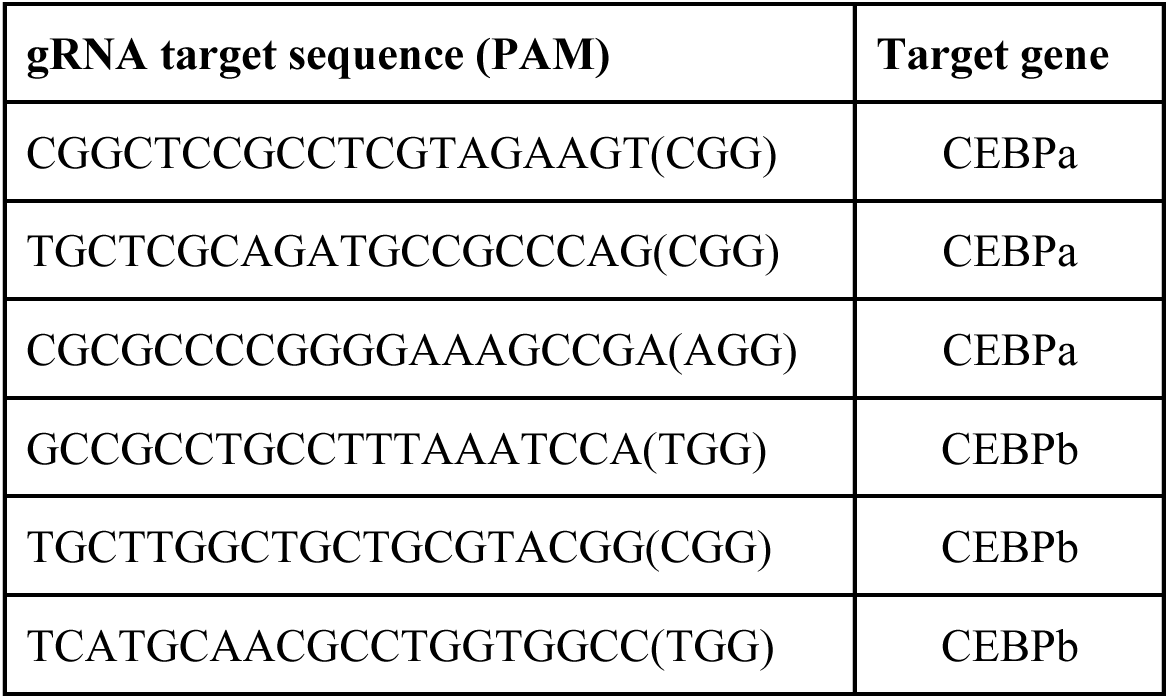

### Genotyping of amplified clonal lines were done by extracting genomic DNA

For genotyping, genomic DNA was extracted from 96-well plates using the DNeasy kit (Qiagen #69504) according to the manufacturer’s protocol. Screening was conducted via colony PCR using the Taq PCR Kit (Qiagen #203203). The presence of *CEBPa/CEBPb* knockouts was assessed using forward and reverse primers designed to anneal approximately 200–400 bp upstream and downstream of the targeted sequence. PCR products were resolved on 1% agarose gel, and KO clones exhibiting either the absence of a PCR product or an altered fragment size were selected for sequencing.

Primers used for KO Screening:

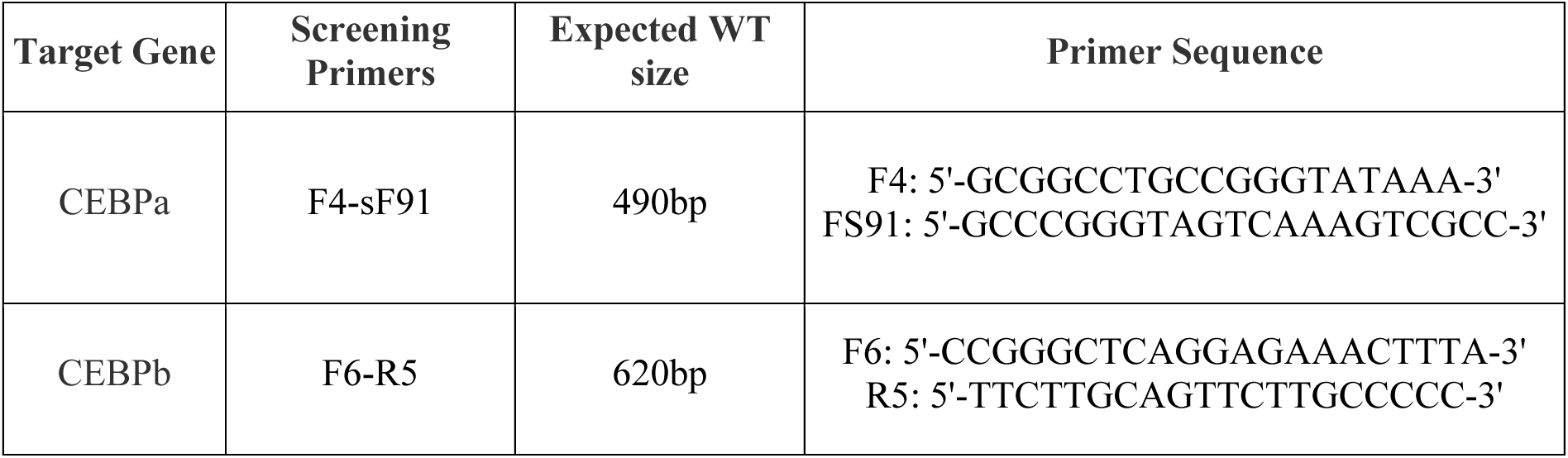
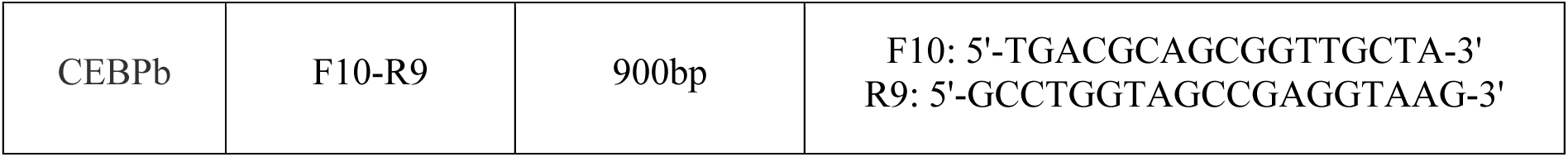

### 10x Genomics Chromium GEX (gene expression) and ATAC assay for transposase-accessible chromatin) library preparation and sequencing

For the scRNA-seq experiments, cells were loaded according to the manufacturer’s protocol for the Chromium Next GEM Single Cell 5’ v2 (DUAL) Kit (10x Genomics) to attain between 500 and 10,000 cells per reaction. Libraries were sequenced, aiming at a minimum coverage of 50,000 raw reads per cell, on the Novaseq 6000 system, using the sequencing format: read 1: 28 cycles; i7 index: 8 cycles, i5 index: 0 cycles, Read 2: 98 cycles. For cell hashing workflow, Chromium Next GEM Single Cell 5′ v2 with Feature Barcoding technology for Cell Surface Protein was used (5’ Feature Barcode kit) according to the manufacturer’s protocol (10x Genomics). The BCR and TCR were sequenced at a target depth of 5,000 reads per cell and following the same sequencing format as the scRNA-seq.

For the scATAC-seq and multimodal snRNA-seq/scATAC-seq experiments, cells were loaded according to manufacturer’s protocol for the Chromium Single Cell ATAC v2 and Chromium Next GEM Single Cell Multiome ATAC+Gene Expression (10x Genomics) to attain between 1,000 and 10,000 nuclei. Libraries were sequencing, aiming at a minimum coverage of 50,000 raw reads per nuclei on the Novaseq 6000 systems, using the sequencing format ATAC v2: read 1: 50 cycles; i7 index: 8 cycles, i5 index: 16 cycles, read 2: 50 cycles. ATAC from multiome: read 1: 50 cycles; i7 index: 8 cycles, i5 index: 24 cycles, read 2: 49 cycles.

### External human bone marrow scRNA-seq and CITE-seq

We collected raw sequencing data from previously published human fetal and adult bone marrow scRNA-seq and CITE-seq datasets. Specifically, we downloaded publicly available .fastq files either from the Gene Expression Omnibus (GEO), ArrayExpress or the European Genome-phenome Archive (EGA). These datasets included: (1) Jardine et al., (E-MTAB-9389, GEO: GSE166895)^24^, (2) Bandyopadhyay et al., (GEO: GSE253355)^29^. We also collected scRNA-seq data of ALL from: (1) Khabirova et al., (EGA: EGAD00001007854)^23^, (2) Caron et al., (GEO: GSE132509)^107^.

### External human fetal YS, AGM, LV, *in vitro* scRNA-seq and CITE-seq

We collected raw sequencing data from previously published human fetal yolk sac(YS), aorta-gonad mesonephros (AGM), liver (LV) scRNA-seq and CITE-seq datasets. Specifically, we downloaded publicly available .fastq files either from the Gene Expression Omnibus (GEO) or ArrayExpress. These datasets included: (1) Goh et al., (E-MTAB-10552, E-MTAB-11549, E-MTAB-11613, E-MTAB-11618)^83–85^, (2) Wang et al., (GEO: GSE144024)^82^, (3) Calvanese et al., (GEO: GSE162950)^46^, (4) Popescu et al., (E-MTAB-7407) ^81^, (5) Suo et al., (E-MTAB-11343)^32^, (6) Alsinet et al., (E-MTAB-11623)^108^.

### Alignment and quantification of scRNA-seq data

Reads from both the newly generated scRNA-seq libraries and external datasets were alignment to the 10x Genomics’ human reference genome GRCh38 (v.3.0.0), followed by cell calling, transcript quantification and QC using the Cell Ranger Software (v.3.0.2; 10x Genomics) with default parameters. *In vitro* datasets cultured with mouse stromal cells were aligned to both GRCh38 (v.3.0.0) and mm10 (v.3.1.0). Cell Ranger filtered count matrices were used for downstream analysis.

### Downstream scRNA-seq and scATAC-seq analysis

#### Donor demultiplexing and doublet identification

We used libraries from the public datasets (E-MTAB-11549, E-MTAB-11613, E-MTAB-11618, GSE166895) which were generated with multiplexed donors. Similarly, for the newly generated libraries, we also multiplexed cell suspensions from multiple donors. To ensure accurate assignment of cells to their respective donor, we genotyped some donors and then pooled sample combinations in a way that each scRNA-seq library contained at least one genotyped donor. To assign each cell in the scRNA-seq libraries back to its donor-of-origin, we first called the single nucleotide polymorphisms (SNPs) in the reads of each barcode with Souporcell^109^ (v.2.4). scRNA-seq reads were genotyped from the Cell Ranger BAM files, aligned to the GRCh38 human reference genome, and compared against the filtered_2p_1kgenomes_GRCh38.vcf reference. Once the cells in scRNA-seq libraries were genotyped, we linked them back to their donor-of-origin genotype using shared_samples.py script from Souporcell across all samples. In yolk sac (YS) and liver (LV), we used the genotype information provided by the original study^83^. Genotype doublets and unassigned cells were discarded in downstream analysis.

### Doublet detection based on transcriptional mixtures

We quantified the likelihood of each barcode to be a cell-doublet with Scrublet software^110^ (v.0.2.3) on a per-library basis. As described in^81^, we performed a two-step diffusion doublet approach to propagate Scrublet scores to similar barcodes, followed by Bonferroni false discovery rate correction. Barcodes estimated as potential doublets were not excluded, but instead these were kept in the downstream analysis and used to identify doublet-enriched clusters after scVI integration and cluster identification (see below).

### Quality filters, batch correction and clustering

We used the filtered count matrices from Cell Ranger 3.0.2 for downstream analysis of the scRNA-seq libraries with Scanpy (v.1.8.2), following their recommended standard practices^111^. We applied stringent QC to further filter the cells called by Cell Ranger to retain only high-quality cells. Specifically, we excluded cells either (1) expressing fewer than 500 genes or (2) with a mitochondrial content higher than 20% or (3) expressing more than 100k UMI counts. For some datasets, our filters discarded more than 50% of the initial called cells.

Next, we flagged cell cycle genes using a data-driven approach as described in^81,112^. To do so, after converting the expression space to log(CPM/100 + 1), where CPM is counts per million, we transpose the object to gene space, performing principal component analysis (PCA), neighbor identification and Leiden clustering. The gene members of the gene cluster encompassing well-known cycling genes (*CDK1, MKI67, CCNB2* and *PCNA*) were all flagged as cell cycling genes, and discarded in each downstream analysis. In parallel, we also used the scanpy function ‘score_genes_cell_cycle’ to infer the cell cycle stage of each cell (that is, G1, G2/M or S) that was later used to interpret the clusters.

Next, we generated an integrated manifold for each study datasets separately. The scRNA-seq manifold included data from two previously published studies as well as the scRNA-seq data newly generated by us. The scVI^113^ (v.0.17.3) low dimensional space was estimated on the top 5,000 most highly variable genes in each dataset. To minimize cell cycle bias, the previously flagged cell cycle genes were excluded before estimating the highly variable genes. We also binned the S and G2/M cell cycle scores (n=20 bins) and used these as covariates for scVI. All the remaining parameters were kept as default, with n_latent =20, n_layers = 2. With the resulting scVI-corrected latent representation of each cell, we estimated the neighbor graph, generated a uniform manifold approximation and projection (UMAP) visualization and performed Leiden clustering. The resolution of the clustering was adjusted manually. The same integration strategy described in the paragraph above was used to reanalyse each of the seven main cell lineages (that is, B cells, T cells, progenitors, myeloid, erythroid, stromal and endothelial cells) to further resolve the cellular heterogeneity in those compartments. Here, we subset the cells to those in the lineage and repeated integration with harmony^114^ (v.0.0.6) or scVI integration using the top 5,000 most highly variable genes within each lineage. The donor and the study ID were kept as batches, with default parameters, adjusting different n_latent and n_layers in each lineage. The resulting harmony or scVI-corrected latent representations were used to derive per-lineage UMAPs and perform Leiden clustering. To create a fetal hematopoiesis integrated manifold, we utilised inVAE^115^ (v.0.1.1) with invariant covariates as broad annotation and organ with 20 dimensions while spurious covariates as sample and sequencing with 5 dimensions. After that, we followed the same downstream procedure as scVI with inVAE latent representation. To create a fetal HSC integrated manifold, we subsetted HSCs cells and followed Scanpy pipeline. And then, we utilised Harmony (v.0.0.6) with 3 principal components, with default parameters, and donor, sequencing type, and cell cycle bins as covariates.

The datasets of human fetal yolk sac, aorta-gonad mesonephros, liver and fetal and adult bone marrow (see section “External human bone marrow scRNA-seq and CITE-seq” and “External human bone marrow scRNA-seq and CITE-seq”) were integrated together with the hiPSC-differentiation cells(Alsinet et al.,^108^ and our hiPSC-B cells differentiation) using scVI, with the most 5,000 highly variable genes and considering as batch the study together with experimental model (*in vitro, in vivo* or cultured fetal bone marrow), the origin (fetal or adult), the organ (yolk sac, aorta-gonad mesonephros, liver and bone marrow), the sequencing type (5’ GEX or 3’GEX) and sequencing platform (scRNA-seq or CITE-seq). The integration was performed using 3 layers and 20 latent variables, followed by Leiden clustering to identify the main clusters and subcluster of progenitors and B-cells. Next, we subset the fetal bone marrow, adult bone marrow, hiPSC-B cells and cultured fetal bone marrow from these two progenitors and B-cells clusters and repeated the scVI integration on the top 5,000 most highly variable genes. Again, to minimize cell cycle bias, the previously flagged cell cycle genes were excluded from the highly variable genes. The integration was performed using 3 layers and 20 latent variables, followed by Leiden clustering and fine-grain annotation of the populations.

### hiPSC-differentiation demultiplexing

First, we used HashSolo^116^ to demultiplex human and mouse cells with a default prior distribution. To distinguish each condition in the *in vitro* experiment, we used HashSolo with a modified prior distribution per each sample. In general, we set higher priors for both negative classification and doublets. Cells assigned as ‘doublet’ or ‘negative’ were excluded from downstream analysis. To verify the above classification, we derived a Gaussian Mixture Model (GMM) with scikit-learn (v0.24) and identified two clusters per each hashtag oligo. From a cluster that has higher hashtag oligo counts, we set a minimum count based on the lower 2nd percentile, and cells with counts below this threshold were also excluded.

### Annotation of cell types

We performed a full re-annotation of the cell clusters in each integrated scRNA-seq manifold generated. When assigning lineage/coarse- and cell type/fine-grained annotations to the Leiden clusters, we first performed a QC round to exclude clusters likely driven by technical artefacts (that is, low QC cells or doublets). Briefly, we flagged as low QC those clusters that (1) express an overall lower number of genes, or (2) express an overall lower number of counts, or (3) display a higher than average mitochondrial RNA content and, importantly, and (4) do not express any distinctive marker gene (and thus are not representing any independent biological entity). Next, we flagged as doublets those clusters that met the following criteria: (1) exhibit higher scrublet doublet score; (2) express marker genes from multiple lineages (for example, display both epithelial and immune markers) and (3) lacked unique gene markers beyond the doublet’s combined gene expression profile. Distinctive marker genes were identified using the Term Frequency–Inverse Document Frequency approach (TF-IDF), as implemented in the SoupX package^117^ (v.1.5.0).

Next, we assigned cell type labels to remaining high-quality clusters. General lineage/coarse annotation was done on the main manifold. Cell state/fine grained annotation was defined on the per-lineage manifold (that is, from reanalyzing the cells in each lineage, as described in the previous section). To aid cell type annotation we also used CellTypist^118^ (v.1.3), a logistic regression classifier optimized by the stochastic gradient descent algorithm. We trained the model by: (1) using both the Pan_Fetal_Human models built into cell types and (2) our own model on the immune cells from the fetal hematopoiesis atlas, this last to annotate the *in vitro* and leukemia datasets. From Jardine and colleagues dataset^24^, we also relied on previous annotations from the author. After projecting inferred labels, we refined the annotations by comparing the expression of distinctive genes (identified with TF-IDF) with bona fide markers from the literature. From the multilineage integration of bone marrow data with published datasets, only cell states with more than 100 cells in the fine-grained annotation were considered for downstream analysis. From the B cell differentiation integration analysis, only fine-grain annotation cell states with more than 150 cells were considered for downstream analysis.

To annotate the leukemia dataset^23^, we followed a workflow similar to that used for the previous datasets. We first annotated cell types using CellTypist trained from our prebuilt fetal and adult bone marrow atlas. We also used the Pan_Fetal_Human models from CellTypist and manually adjusted cell annotations based on comparisons with our fetal and adult bone marrow atlases.

### Integration our fetal bone marrow dataset with an inDrop-based scRNA-seq dataset

To annotate published bone marrow dataset from inDrops technology^30^, we downloaded the count matrices and integrated our generated fetal bone marrow dataset. After applying standard quality control (removing cells expressing fewer than 500 genes), we used scVI^113^ (v.1.1.15) to estimate a low dimensional space based on the top 3,000 most highly variable genes. The donor was treated as a batch, and the study was added as a covariate. All the other parameters were kept as default, with n_latent =10, n_layers = 1. Using the resulting scVI-corrected latent representation of each cell, we estimated the neighbor graph, generated a uniform manifold approximation and projection (UMAP) visualisation, and annotated cell types using a CellTypist trained on our prebuilt fetal and adult bone marrow atlas.

### BCR/TCR

All single-cell V(D)J data from the 5′ Chromium 10X kit were initially processed with cellranger vdj pipeline (v.6.1.2) with cellranger vdj GRCh38 reference (v.5.0.0). BCR contigs contained in ‘all_contigs.fasta’ and ‘all_contig_annotations.csv’ were then processed further using dandelion singularity container^48^ (v.0.3.7). Specifically, V(D)J output data were re-annotated with igblastn against IMGT (international ImMunoGeneTics) reference sequences by Dandelion preprocessing. For *in vitro* BCR data, we skipped tigger which reassigned heavy chain V gene alleles to avoid errors occurring from low quality contigs. From Dandelion output, we used ‘all_contig_dandelion.tsv’ output, while we used ‘all_contig_igblast_db-all.tsv’ to analyze productive and nonproductive repertoires. In general, we exclude cell types if they were less than 100 cells from each study. To define productive and nonproductive, it was defined by True and False from the productive_ratio function for IGH, IGK, and IGL respectively. For most repertoire characteristics including isotype, mutation frequency, junction length, np length and gene usage, we used Dandelion output and were analyzed per donor. In gene usage, we performed multiple testing corrections with the FDR approach using p.adjust function from stats (v.4.0.4). In gene usage, we ignored polymorphic variants from IMGT gene names to reduce the data groups and downsampled per donor to match the same number of contigs. For the visualization, we removed genes that were less than 1% in average and re-ordered based on genomic position from IMGT. In isotype, we remove multi_contig from the analysis. To analyze repertoire diversity metrics, we used Immunarch^119^ (v.0.9.1). For the input of the data, we first subsetted per cell type and per donor and downsampled to have the same number of contigs per donor from Dandelion output. To identify overlapping repertoire, we measured Morisita index using the repOverlap function with method = ‘morisita’. To investigate repertoire diversity, we measured Inverse Simpson index using the repDiversity function with method = ‘inv.simp’. To analyze gene usage overlap, we measured jensen-shannon using geneUsage function, following geneUsageAnalysis function with method = ‘js’. For all statistical difference tests, the glm function from stats (v.4.0.4) was used.

### Differential gene expression fetal vs adult bone marrow

We evaluated the magnitude and significance of gene expression differences between subpopulations across the B cell lineage or the stromal cells in fetal and adult bone marrow using limma^120^ (v.3.46.0). First, in case of the B cell lineage, we downsample to 500 cells per subpopulation. Secondly, to account for within-sample correlations (i.e. cells coming from the same donor), pseudobulking with sum aggregation was performed prior to applying limma. Specifically, we generated three pseudobulks per donor and per cell type by aggregating the cells of each cell type and taking the mean gene expression within the cell type. Finally, we tested for differential expression between conditions (fetal vs adult) using the “limma-voom” approach. The analysis was performed on the scRNA-seq datasets, and we reported as differentially expressed genes with logFC > 0 and an adjusted p-value < 0.05.

### Cell–cell communication analysis with CellPhoneDB

We used CellPhoneDB v5.0.0^58^ to study cell-cell interactions between the bone marrow microenvironment and B cell lineage cells, identifying those differentially expressed between fetal and adult. Using CellPhoneDB method-3 based on differential gene expression, we retrieved pairs of interacting ligands and receptors meeting the following requirements: (1) all the interacting partners were expressed by at least 5% of the cell type under consideration; (2i) the receptors were significantly overexpressed by the B cells; or (2ii) the ligands were significantly overexpressed by the bone marrow microenvironment. Differential expression analysis was performed on a per-cell type basis comparing matched fetal and adult cell types along the B-cell lineage trajectory (from HSC to preNaiveB) or in the bone marrow microenvironment using the approach described above.

### Trajectory inference and differential expression along trajectories

We used the trajectory inference method Slingshot^121^ (v1.8.0) to model B cell differentiation in fetal and adult bone marrow, designating hematopoietic stem cells (HSC) as the starting point of the trajectory and preNaiveB as the endpoint. Each pseudotime was then scaled between 0 and 1. The scaled pseudotime ordering of the cells along with the weighted assignment was then used as input for TradeSeq^122^ (v1.18.0) to extract genes that are differentially expressed along the B-cell lineage or between fetal/adult with the *associationTest()* function.

### Differential gene expression leukemia vs adult bone marrow

To identify the genes differentially expressed in the ELP/CLP subpopulation between each leukemia patient compared to adult bone marrow, we used the *sc.tl.rank_genes_groups* function from scanpy (v1.8.2) using the two-sided Wilcoxon rank-sum test. Genes with log fold-change (logFC) > 1 and adjusted p-value < 0.01 were considered upregulated in leukemia.

Subsequently, we compared these leukemia upregulated genes to those upregulated in fetal versus adult bone marrow. This allowed us to extract the genes that were consistently upregulated in both leukemia and fetal bone marrow cells compared to their adult counterparts.

### Chromatin accessibility data preprocessing

For single-cell ATAC-seq, we applied ArchR^123^ (v1.0.2) to process the outputs from CellRanger-ARC. Initial per-droplet quality control was performed considering the number of unique nuclear fragments, signal-to-background ratio and the fragment size distribution. Moreover, droplets with transcription start site enrichment score < 4 and number of fragments < 1,000 were removed. Doublets were discarded using the default settings. Initial clustering was performed at a resolution of 0.8 on the embedding from iterative latent semantic indexing using default values. Then, pseudo-bulk replicates were made for each broad cluster per region from the initial clustering results. Peak calling (501-bp fixed-width peaks) was performed based on pseudo-bulk coverages by MACS2 (v2.2.7.1). Then, a cell-by-peak count matrix was obtained and exported. Peaks were extracted and integrated with RNA count matrix data using GLUE^124^. Cell type labels were transferred from the RNA to the ATAC data via the shared knn-graph using majority voting. Peak calling was then repeated per cell type and new cell-by-peak count matrices were created before and after subsetting to the B cell lineage, starting from hematopoietic stem cells to preNaiveB cells. The peak locations reported by ArchR were used to summarise the peak types (promoter/exonic/intronic/distal/). In addition, the peak matrices for fetal and adult bone marrow were used to analyse the conserved and specific peaks, selecting the conserved peaks found in 90% of the cells and specific peaks below 10% of the cells. Only fine-grained cell states with equal or more than 20 nuclei were considered.

### Gene regulatory network (GRN) analysis

The SCENIC+^57^ (v1.0.0) pipeline was used to predict transcription factors and putative target genes as well as regulatory genomic regions with binding sites. A common matrix with consensus peaks called using ArchR for prenatal and adult bone marrow was built using the iterative overlap peak merging procedure^125^. We then subsetted for the B cell differentiation trajectory from hematopoietic stem cells (HSC) to preNaiveB cells for downstream analyses. Pseudo-multiome meta-cells were created from separate RNA and ATAC by clustering droplets into groups of around 10–15 droplets based on a bipartite knn-graph constructed from the shared RNA and ATAC embedding, and then aggregating counts and fragments subsequently. The pipeline was applied to subsets of the dataset corresponding to individual lineages: First, CisTopic (pycistopic v1.0.2) was applied to identify region topics and differentially accessible regions from the fragment counts as candidate regions for transcription factor binding. CisTarget (pycistarget v1.0.2) was then run to scan the regions for transcription factor-binding sites, and GRNBoost2 (arboreto v0.1.6)^126^ was used to link transcription factors and regions to target genes based on co-expression or accessibility. Enriched transcription factor motifs in the regions linked to target genes were used to construct transcription factor–region and transcription factor–gene regulons. Finally, regulon activity scores were computed with AUCell based on target gene expression and target region accessibility, and regulon specificity scores derived from them. Networks of transcription factors, regions and target genes (enhancer-driven GRNs) were constructed by linking individual regulons.

### SNP2Cell

The SNP2Cell tool^127^ was applied to compute SNP scores for the genes and distal regulatory elements forming the gene regulatory networks of B cell development. Briefly, SNP scores were computed from published full GWAS summary statistics and weighted by linkage disequilibrium. SNP scores were then mapped to regulatory elements within the gene regulatory networks, and propagated to nearby TFs and genes in the network^128^. Statistical significance was determined based on random simulations of perturbations. Analogously, marker gene scores for each cell cluster were mapped to the network and propagated to detect cell cluster specific network nodes. Finally, both types of scores were combined to identify cell-type-specific disease-associated sub-networks.

### Pseudotime estimation from joint embedding of *in vitro*, cultured fetal bone marrow and *in viv*o

From *in vitro* (hiPSC-differentiation dataset), cultured fetal bone marrow dataset and *in vivo* bone marrow (Vilarrasa-Blasi dataset), we extracted cell types representing the B cell developmental trajectory (that is from HSC, MPP, LMPP, ELP/CLP, PreProB, EarlyProB, LateProB, LargePreB, SmallPreB, ImmatureB, PreNaiveB, B1, MemoryB and Plasma). To generate an integrated manifold, we followed the same procedure previously applied to create the *in vivo* bone marrow manifold. Here, we used as a batch the combination of study, experimental models (*in vitro, in vivo* or cultured fetal bone marrow), origin (fetal or adult), organ (yolk sac, aorta-gonad mesonephros, liver and bone marrow), sequencing type (5’ GEX or 3’GEX), and sequencing platform (scRNA-seq or CITE-seq) together with binned cell cycle scores as covariate, as previously described. All the remaining parameters were kept as default, with n_latent =20, n_layers = 2. The resulting scVI-corrected latent representation was used for Leiden clustering. To reduce computational demands for downstream trajectory inference analysis, we created a reduced representation of the manifold while preserving their underlying cell-type composition. This was achieved by randomly subsampling each cell type to 100 cells for each *in vitro*, cultured fetal bone marrow, fetal *in vivo* bone marrow and adult *in vivo* bone marrow (if the cell type had less than 100 cells, the original cell number was kept). From this reduced embedding, we estimated the differentiation pseudotime from HSC to immatureB by employing Slingshot^121^ (v.1.8). From Slingshot, we ran default parameters with thresh = 1 and did not specify start and end clusters. At the end, we scaled the resulting pseudotime from 0 to 1 for each experiment.

### *In vivo* and *in vitro* trajectory alignment analysis by Genes2Genes

To evaluate the agreement between *in vivo* (fetal and adult bone marrow) and *in vitro* (hiPSC-differentiation) B-cell differentiation trajectories, we utilized Genes2Genes^89^ (v.0.2). This Bayesian Information-theoretic Dynamic Programming framework is designed to effectively capture matches and mismatches between two trajectories. Before alignment, both the *in vitro* and *in vivo* B cell trajectories were discretised over the pseudotime axis from Slingshot resulting in equal length intervals. To discretise the time points, we inferred the optimal binning of the pseudotime distribution using the ContinuousOptimalBinning function from optBinning^129^ (v.0.20.0), which determined the number of discrete pseudotime points as 15. After binning, we identified that 2% of the adult ELP/CLP cells were misassigned to the last pseudotime point and were discarded. We then examined the trajectory alignment, focusing on the dynamics of human TFs (n=1,555) and other genes associated with other genes associated with B-cell biology (n=279). We also performed hierarchical clustering on the gene space to identify groups of genes that show similar temporal alignment. The number of gene clusters was decided based on a distance threshold that showed a good tradeoff between the number of clusters versus mean silhouette coefficient. From the clustering results, we checked the genes of the cluster with the highest opt_alignment_cost and the lowest alignment_similarity_percentage. We also selected the top mismatching genes (with ≤ 50% alignment similarity) between *in vivo* and *in vitro* and studied their enrichment in signaling pathways using GSEApy wrapper against the MSigDB_Hallmark_2020 and KEGG_2021_Human pathway gene sets.

### Human orthologue genes for mouse stromal

Mouse stromal cells were filtered using the parameters outlined in the quality filters section. Genes expressed in mouse stromal cells were matched to their human orthologues using the ortholog_one2one function in Ensembl. The resulting expression matrix of mouse cells with human orthologous genes was then combined with the human cells in the hiPSC-differentiated dataset to identify ligands specifically produced by the mouse cells (with cognate receptors present in our hiPSC-differentiated subpopulations, according to CellPhoneDB database).

